# Comparative analysis of commercially available single-cell RNA sequencing platforms for their performance in complex human tissues

**DOI:** 10.1101/541433

**Authors:** Yue J. Wang, Jonathan Schug, Jerome Lin, Zhiping Wang, Andrew Kossenkov, the HPAP Consortium, Klaus H. Kaestner

**Affiliations:** Department of Genetics and Institute for Diabetes, Obesity, and Metabolism, University of Pennsylvania Perelman School of Medicine, Philadelphia, PA; Department of Biomedical Sciences, Florida State University College of Medicine, Tallahassee, FL, USA; Bioinformatics core, Institute for Biomedical Informatics, University of Pennsylvania Perelman School of Medicine, Philadelphia, PA; Bioinformatics facility, The Wistar Institute, Philadelphia, PA; The Human Pancreas Analysis Program (RRID:SCR_016202)

## Abstract

The past five years have witnessed a tremendous growth of single-cell RNA-seq methodologies. Currently, there are three major commercial platforms for single-cell RNA-seq: Fluidigm C1, Clontech iCell8 (formerly Wafergen) and 10x Genomics Chromium. Here, we provide a systematic comparison of the throughput, sensitivity, cost and other performance statistics for these three platforms using single cells from primary human islets. The primary human islets represent a complex biological system where multiple cell types coexist, with varying cellular abundance, diverse transcriptomic profiles and differing total RNA contents. We apply standard pipelines optimized for each system to derive gene expression matrices. We further evaluate the performance of each system by benchmarking single-cell data with bulk RNA-seq data from sorted cell fractions. Our analyses can be generalized to a variety of complex biological systems and serve as a guide to newcomers to the field of single-cell RNA-seq when selecting platforms.

## INTRODUCTION

Single-cell RNA-seq technologies offer an unprecedented precision in dissecting the molecular phenotypes of complex biological systems. Currently, there are three major commercially available single-cell RNA-seq platforms: Fluidigm C1, Clontech iCell8, and 10x Genomics Chromium. These platforms differ in their sensitivity, specificity, throughput, and other experimental parameters.

Previously, several groups performed comparisons between different single-cell RNA-seq platforms and chemistries (Svensson et al., 2017; Torre et al., 2018; Wu et al., 2014; Zhang et al., 2019; Ziegenhain et al., 2017). However, these analyses either used clonal cell lines with rather uniform expression profiles or synthesized spike-in RNAs, and thus do not recapitulate the complexity of the transcriptomes from heterogeneous biological system (Risso et al., 2014). In addition, these comparisons utilized home-build drop-seq platforms and/or customized reaction chemistries, which can potentially be a hurdle for investigators new to the single-cell RNA-seq field. Here, we systematically compared three of the most popular commercial single-cell platforms using primary pancreatic islets cells from human donors. The human pancreatic islet contains at least five types of endocrine cells, together with endothelial cells, mesenchymal cells, exocrine cells and other rare cell types. As such, the pancreatic islet represents a complex biosystem bearing the characteristics of other primary organs and tissues. Therefore, it serves as an ideal system to evaluate the performance of different RNA-seq platforms. Here, we report the sensitivity, accuracy, doublet rate, throughput and cost of the three most popular commercial single-cell platforms, using standard protocols provided by manufacturers. Our study provides essential performance parameters of different platforms that can be critically evaluated by individual investigator to tailor their research needs.

## RESULTS

### Overview of commercial single-cell RNA-seq platforms

The three commercial single-cell RNA-seq platforms are highly divergent in their approaches and chemistries. In the Fluidigm C1 platform, the integrated microfluidic chip (IFC) is utilized as a reservoir to capture, image, and perform cell lysis, reverse transcription (RT) and initial PCR reactions for single cells. There are two types of IFCs: 96 and 800 HT, which have the capacity to capture up to 96 cells and 800 cells, respectively. In addition, each type of IFCs comes with three different reaction chamber sizes (5-10 μm, 10-17 μm, and 17-25 μm) to facilitate capturing of cells ranging from 5 μm to 25 μm in diameter. The basis of Clontech’s iCell8 system is an alloy nanogrid wafer with 5,184 nanowells. Within each nanowell, oligonucleotides containing poly d(T), unique molecular identifiers (UMIs) and unique well barcodes are preprinted. An integrated robotic system dispenses cells into each of the nanowell from a master 384-well plate. According to Poisson distribution, on average, optimally ~1,800 single cells can be captured. Single live cells can be recognized and selected by the automated imaging station. The iCell8 system can accommodate cells up to 100 μm in diameter (Goldstein et al., 2017). A third singlecell RNA-seq system, the 10x Genomics Chromium platform, is a droplet-based system. Its ‘Gel-bead in Emulsion’ (GEM) technology allows for capturing of an average of 50% of input cells. Each gel bead is coated with oligonucleotides containing RT primers, UMIs and cell barcodes. For a typical Chromium experiment, 10,000 cells are loaded and approximately 5,000 cells can be captured and sequenced. Two versions of the Chromium single-cell RNA-seq chemistry have been released (V1 and V2), with the difference in their final RNA-seq library configuration. During the preparation of this manuscript, 10x Genomics released a V3 chemistry kit, which further improves the single-cell RNA-seq sensitivity and incorporates feature barcoding that allows for the quantification of cell surface proteins (Stoeckius et al. 2017). 10x Genomics recommends processing cells less than 50 μm in diameter in the system (Zheng et al., 2017). Due to the unique designs, the three single-cell platforms differ in throughput, individual cell trackability and final single-cell libraries (Table 1). For example, because of the inclusion of imaging modules, individual cells in the C1 96 and iCell8 platforms and each combination of 40 cells in the C1 HT system can be independently selected for downstream library preparation and/or deeper sequencing. On the other hand, in the Chromium system, the entire library pool is sequenced and the identification of individual cell is performed *post hoc.* In the basic C1 protocol, no UMI is used for library construction, while in the iCell8 and Chromium systems, the usage of UMI allows for absolute counting of transcript numbers (Islam et al., 2014). In addition, multiple samples can be labelled by unique fluorescent markers and combined in one single-cell capture experiment in the C1 and iCell8 platforms, while for Chromium, the recent development of the cell hashing method with DNA-barcoded antibodies allow for sample multiplexing (Stoeckius et al. 2017).

**Table 1.** Properties of single-cell RNA-seq platforms. Related to Figure 1.

|  | C1 96 | C1 HT | iCell8 | Chromium |
| --- | --- | --- | --- | --- |
| Imaging capability | Yes | Yes | Yes, built-in with 4x magnification | No |
| Transcriptomic data | Full length | 3' | 3' | 3' or 5' |
| # of cells passed QC | 1,248/1,305 (96%)<br>(read depth filtering, median -2SD) | 5,058/5,353 (94%)<br>(read depth filtering, median-2SD) | 411/952 (43%) | 17,531/18,213 (96%) |
| Throughput | Up to 96 cells | Up to 800 cells | ~ 1000 cells | ~ 5000 cells |
| Selection of individual cell | Yes | Per row | Yes | No |
| UMI | No (optional) | No (optional) | No (optional) | Yes |
| ERCC spike-in | Yes | Yes | Yes | No |

Human pancreatic islets contain a variety of cell types, including five endocrine cell types (including alpha, beta, delta, epsilon, and PP cells), ductal, acinar, endothelial, as well as fibroblasts cells. Aberrations in the molecular phenotypes of these cells are highly relevant to the pathogenesis of metabolic disorders such as diabetes. Therefore, human pancreatic islets present an ideal system to determine real-world performance parameters of the three single-cell RNA-seq platforms.

### Sequencing quality control and data preprocessing

We collected single cells from one to twenty-five samples depending on the platform (Figure 1 and Table S1) and performed quality control steps to filter out uninformative cells. Due to the differing chemistries and throughput of individual single-cell platform, we applied separate thresholds to each platform. For the C1 96, C1 HT and iCell8 system, we analyzed the microscope images of the capture sites and only selected single cells for further analyses. On average, 42 ± 4 out of the 96 (44%) capture sites on the C1 96 Chip were occupied by a single cell with no visible debris (a total of 1,305 cells from 35 captures). On the C1 HT platform, an average of 329 ± 27 out of the 800 (41%) capture sites were occupied by a single live cell (a total of 5,353 cells from 17 captures). In the one iCell8 run, 952 cells were identified as single live (19%). Only annotated single cells from these three systems were carried into RNA-seq library preparation and further analyzed. For the Chromium systems, since there was no visual guide on the capture and no physical way to pre-select individual cells, all cells were pooled and further processed. To eliminate cells with technical failures, in the C1 96 and C1 HT platforms, we excluded cells with reads lower than 2 standard deviation from the population median. 1,248 (96%) cells from the C1 96 platform and 5,058 (94%) cells from the C1 HT platform passed the read count filter, with an average sequencing depth of 2.2×10^6^ for the former and 4.9×10^5^ for the latter. In the iCell8 system, we used a minimum read depth of 10^5^ as cutoff and 411 (43%) cells remained after filtering, with an average of 3.8×10^5^ reads per cell. For the Chromium systems, we applied 10x Genomics recommended thresholding methods based on Cell Ranger output to identify cells. Briefly, cell barcodes were ordered based on the sum of UMIs associated with each barcode. The threshold for good quality cells were chosen where the UMI counts for all the selected cells were at least 1/10 of the 99^th^ percentile of the UMI counts of the expected recovered cells (Figure S1A). To ensure that we only selected single viable cells for downstream processing, cells were further filtered based on numbers of genes detected with a lower limit of 200 and an upper limit of 4,000 for Chromium V1 and 7,500 for Chromium V2. We further filtered out cells with more than 10% mitochondrial reads, since high proportions of mitochondrial alignments are indicative of impaired intracellular membranes (Ilicic et al. 2016). Based on these filtering strategies, 5,045 cells from Chromium V1 and 18,213 cells from Chromium V2 were carried on for downstream analyses. The average sequencing depth was 7.3×10^4^ reads/cell for the Chromium V1 and 2.1×10^5^ reads/cell for the Chromium V2 chemistry. Sequencing saturation in the Chromium system was calculated as the percentage of non-unique UMIs within the total number of reads. The Chromium V1 library was sequenced to 75% saturation and the Chromium V2 libraries were sequenced to an average of 78% saturation. We plotted the dependency of average number of genes detected with at least 1 read on sequencing depth for all single-cell platforms (Figures S1B and S1C). We observed reasonable sequencing depth in all platforms (Figures S1B and S1C). The thresholds and basic sequencing statistics for different libraries are summarized in Figures S1B-S1C and Table 2.

**Figure 1.**
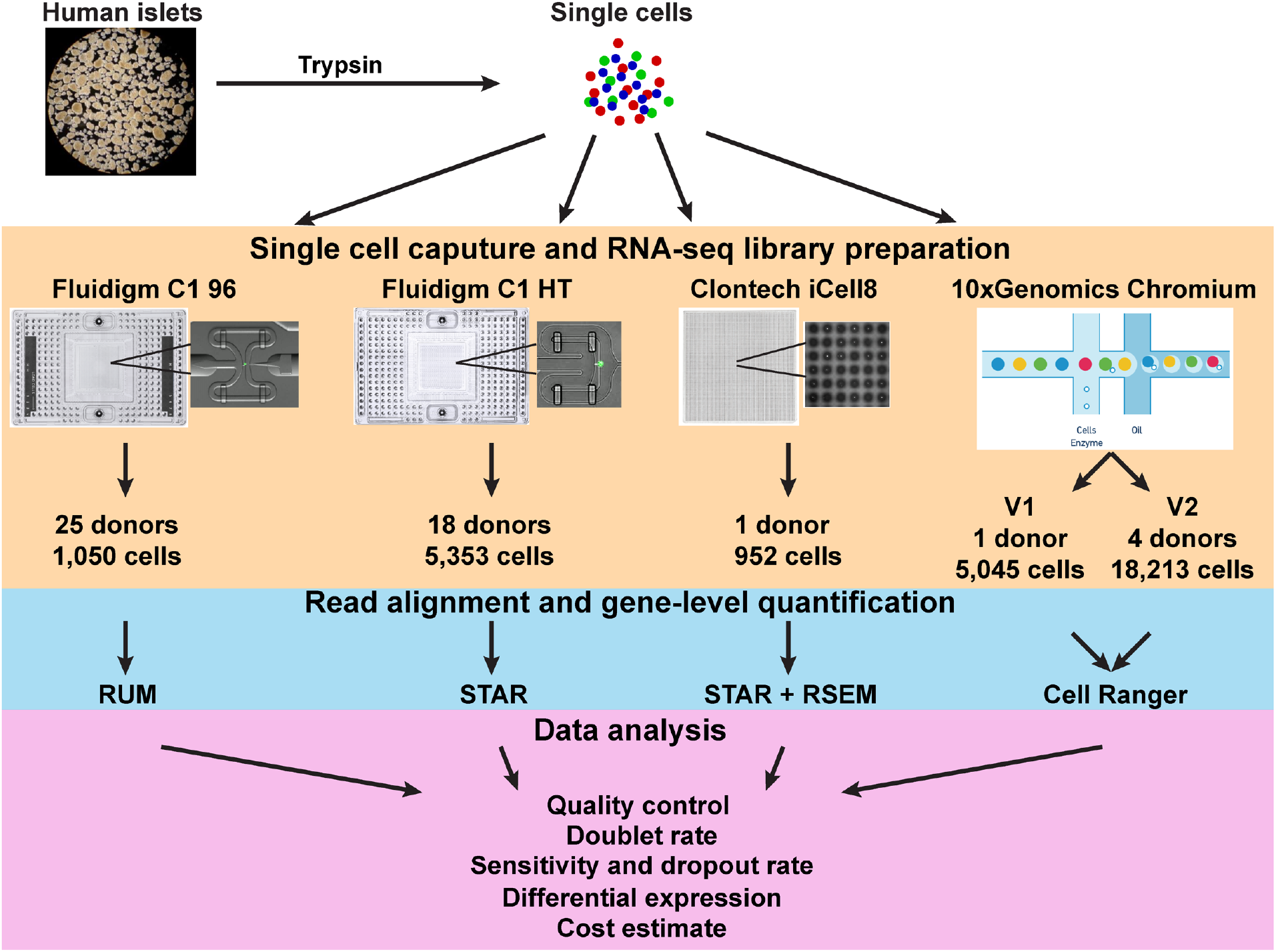
Study overview. See also Figure S1, Tables 1, 2 and S1.

### Doublet rate

To ensure that single-cell transcriptomic data represent true single cells rather than aggregates of two or more cells, the doublet rate is a critical parameter to be evaluated. In the C1 96, C1 HT and iCell8 systems, images of the capture sites facilitate the identification of single cells. However, some ‘stacked doublets’ and small cellular debris may be missed because of the limited resolution of microscopy scanning. In the Chromium system, the overabundance of gel beads relative to cells keeps the doublet rates at a theoretically low level. We leveraged the fact that the predominant pancreatic endocrine cells are single hormone positive to estimate the doublet rate in our single-cell experiment (Wang et al., 2016b; Xin et al., 2016). In particular, we used the percentage of double positivity for INS and GCG, makers of beta and alpha cells, respectively, as a surrogate to estimate the percentage of doublets (Figure 2). We adjusted the observed rate of INS+GCG+ cells by the relative abundance of the INS+ and GCG+ populations (doublet rate = (observed percentage of double positive cells)/(2*(frequency of INS+ cells)*(frequency of GCG+ cells)). We found that the doublet rate was 25% in the C1 96 platform, 33% in the C1 HT platform, 26% in the iCell8 platform, 28% in the Chromium V1 platform and 21% in the Chromium V2 platform. In general, the doublet rates we estimated here are higher than what was previously reported in the different platforms based on mixed species experiment with human and mouse cell lines (Fluidigm; Goldstein et al., 2017; Zheng et al., 2017). This likely reflects the difficulties associated with generating true single cell suspensions from solid tissue using enzymatic digestion, and the propensity of epithelial cells to reaggregate after dissociation. Alternatively, it is possible that the mixed species cell experiment frequently employed to judge doublet rates underestimates the true double rate because mouse and human cell lines are not likely to re-aggregate in suspension. Regardless, for investigators analyzing primary cells other than blood cells, the presence of doublets should be taken into account to avoid spurious conclusions.

**Figure 2.**
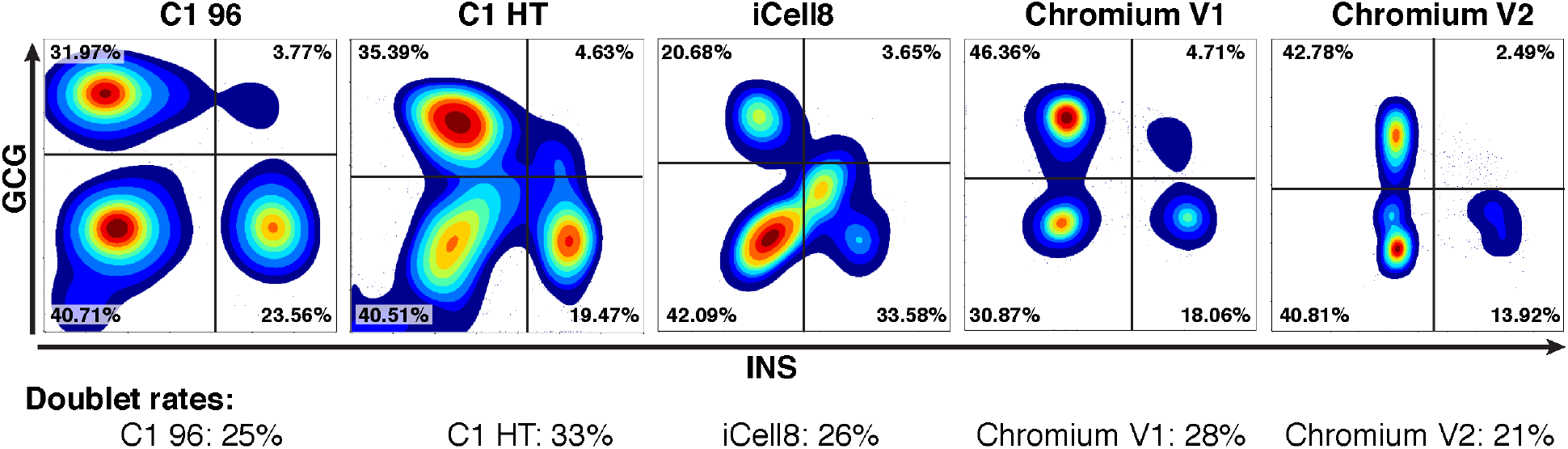
Doublet rate estimates. INS versus GCG density plots are shown for each single-cell platforms. Data for multiple donors are pooled. On the Chromium V2 platform, only data from the two control donors were used due to non-detectable INS expressions in the two T1D donors. The percentage of cells falling within each quadrant is shown on the graph. The calculated doublet rates for each platform are displayed underneath the corresponding graph.

### Detection of individual pancreatic cell types

To evaluate the capability of the three platforms to identify cell type heterogeneity, we employed the pipeline from the Seurat package (Butler et al., 2018). Briefly, we performed principal component analysis (PCA) with the most variable genes, clustered the cells using a graph-based approach, employed tSNE for visualization, and finally annotated cell types based on known marker genes. Cell type identification was performed independently for each platform to avoid potential confounders associated with technical variation. Different cell types largely located to distinct clusters in the tSNE graphs and we could identify major cell types including alpha, beta, acinar, ductal, endothelial and mesenchymal cells in the majority of the platforms (Figure 3). However, unique clusters for delta, epsilon, and PP cells were rarely observed, likely due to their low abundance and transcriptomic similarities to alpha and beta cells. In addition, in the iCell8 system, alpha and beta cells were not readily distinguishable with the basic Seurat pipeline (Figure 3C). To resolve the two different endocrine populations in the iCell8 data, we subsequently utilized single-cell consensus clustering (SC3) with k=8 for hierarchical inference (Kiselev et al., 2017). Alpha cells and beta cells were thereby separated in the SC3 plot based on high expression of GC, PCSK2 and IRX2 for alpha cells and ABCC8 and PCSK1 for beta cells (Figure S2). We projected the cell type labels generated from SC3 algorithm onto the tSNE plot generated from Seurat pipeline and observed that the results from these two pipelines were mostly in agreement with each other (Figure 3C). Finally, on the tSNE plots obtained from the C1 HT and iCell8 platforms there was a group of cells with ambiguous marker gene expression (“other” in Figures 3B and 3C). Since these cells generally had intermediate levels of multiple markers, they may represent doublets, debris, or dying cells.

**Figure 3.**
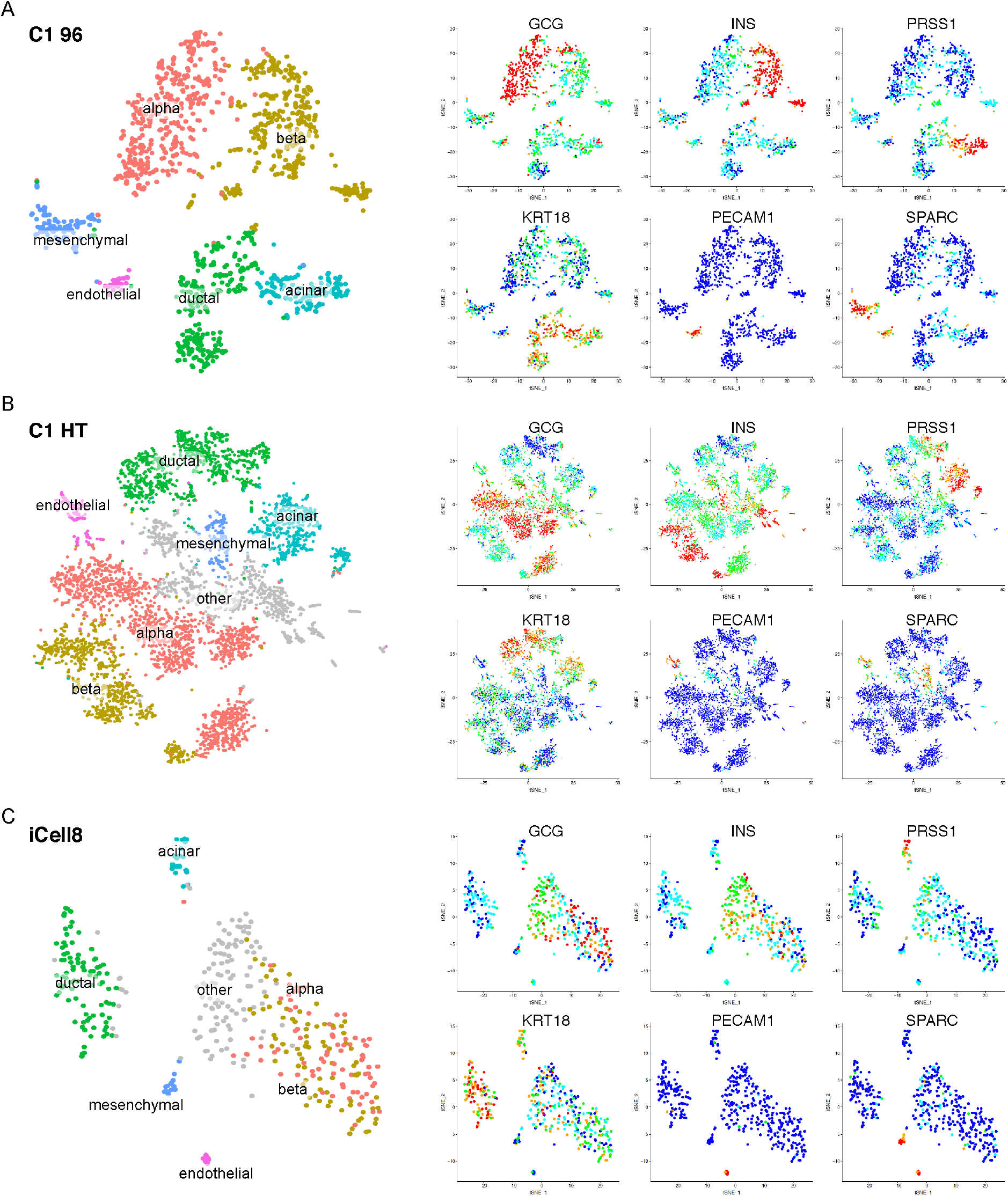
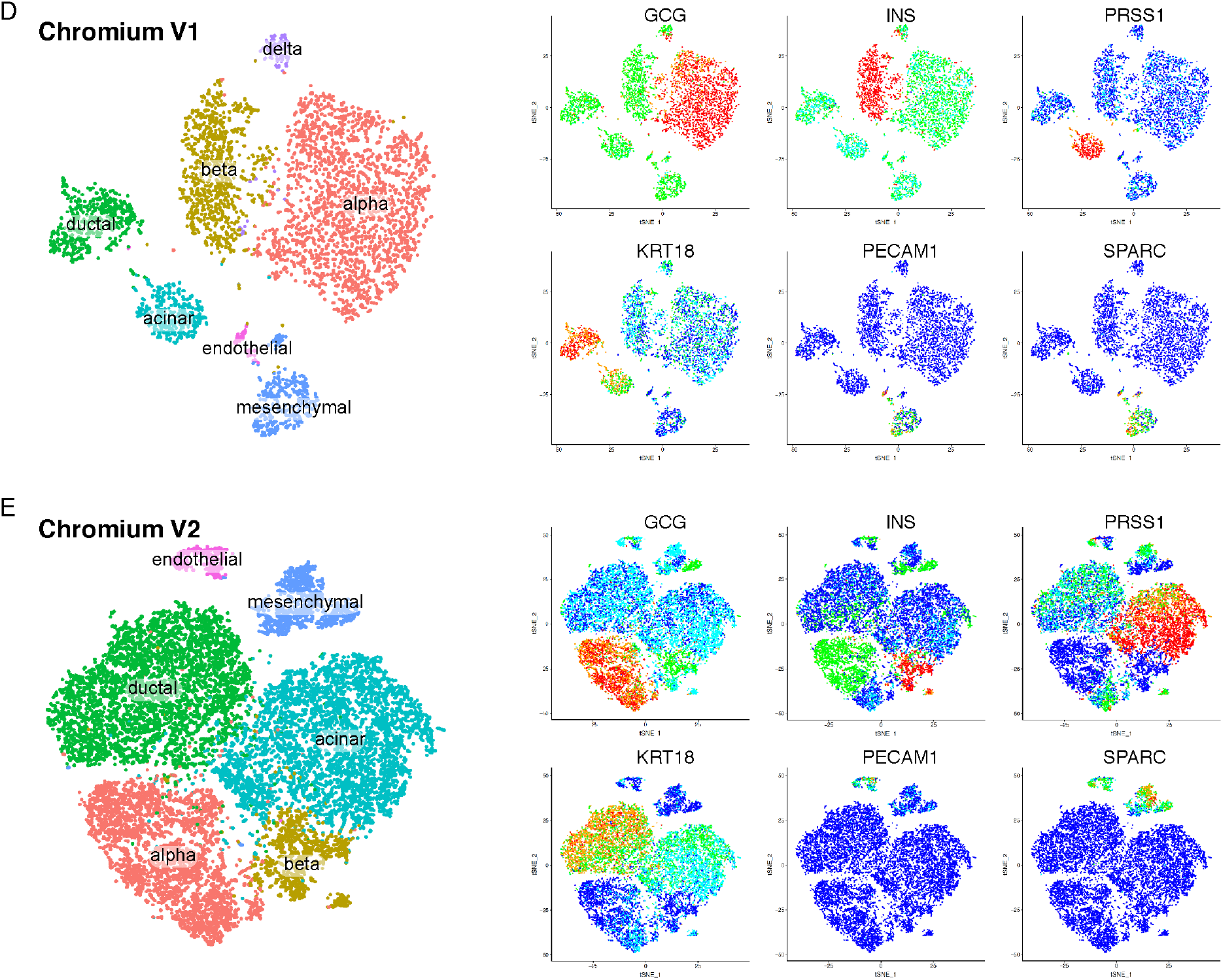
Single-cell RNA-seq can distinguish different pancreatic cell types. See also Figures S2 and S3. tSNE plots are shown for single-cell data generated on (A) C1 96, (B) C1 HT, (C) iCell8, (D) Chromium V1 and (E) Chromium V2 platforms. Cell type labels are shown on the tSNE plot at the left side of each panel. Marker gene expressions are shown on the six tSNE plots at the right side of each panel.

Next, we plotted pairwise relationships of the read counts from different platforms within each cell type. We observed a high correlation between all platforms (Figure S3). For all cell types, the highest correlation was observed between Chromium V1 and Chromium V2 systems, and the lowest correlation was most frequently between C1 96 and Chromium V1 libraries (Figure S3).

### Sensitivity and dropout rate

Because of the technical limitations to capture RNA molecules, especially those with low abundance, not all genes can be detected by current single-cell transcriptomics, and ‘dropout’ events, or zero reads for an mRNA that is normally expressed in a given cell type, are frequently observed (Kharchenko et al., 2014). Because this is an important performance issue, we quantified the number of genes detected as transcribed per cell (reads or UMI ≥ 1) to evaluate the sensitivity of different platforms. The C1 96 platform had the highest sensitivity compared to all the other systems (Figure 4A). Depending on cell types, the median number of genes detected as transcribed per cell ranged from 5,498 to 7,249. The iCell8 and the C1 HT platforms performed similarly, with between 4,414 to 5,450 transcribed genes per cell identified in the former versus 3,899 to 4,880 in the latter. The Chromium V2 system ranked next with between 2,961 to 4,109 genes detected as transcribed, while the Chromium V1 showed the worst performance (1,439 to 2,252). To separate out the effect of sequencing depth, we down-sampled sequencing reads from all non-Chromium V1 platforms to 7×10^4^ reads/cell, a level that matched the sequencing depth in the Chromium V1 platform. Even with the artificially reduced read counts, we again observed that the highest number of genes detected as transcribed per cell in the C1 96 platform, followed by the iCell8 and C1 HT systems (Figure 4B). At this sequencing depth, the sensitivities of the Chromium V1 and Chromium V2 platforms were comparable.

**Figure 4.**
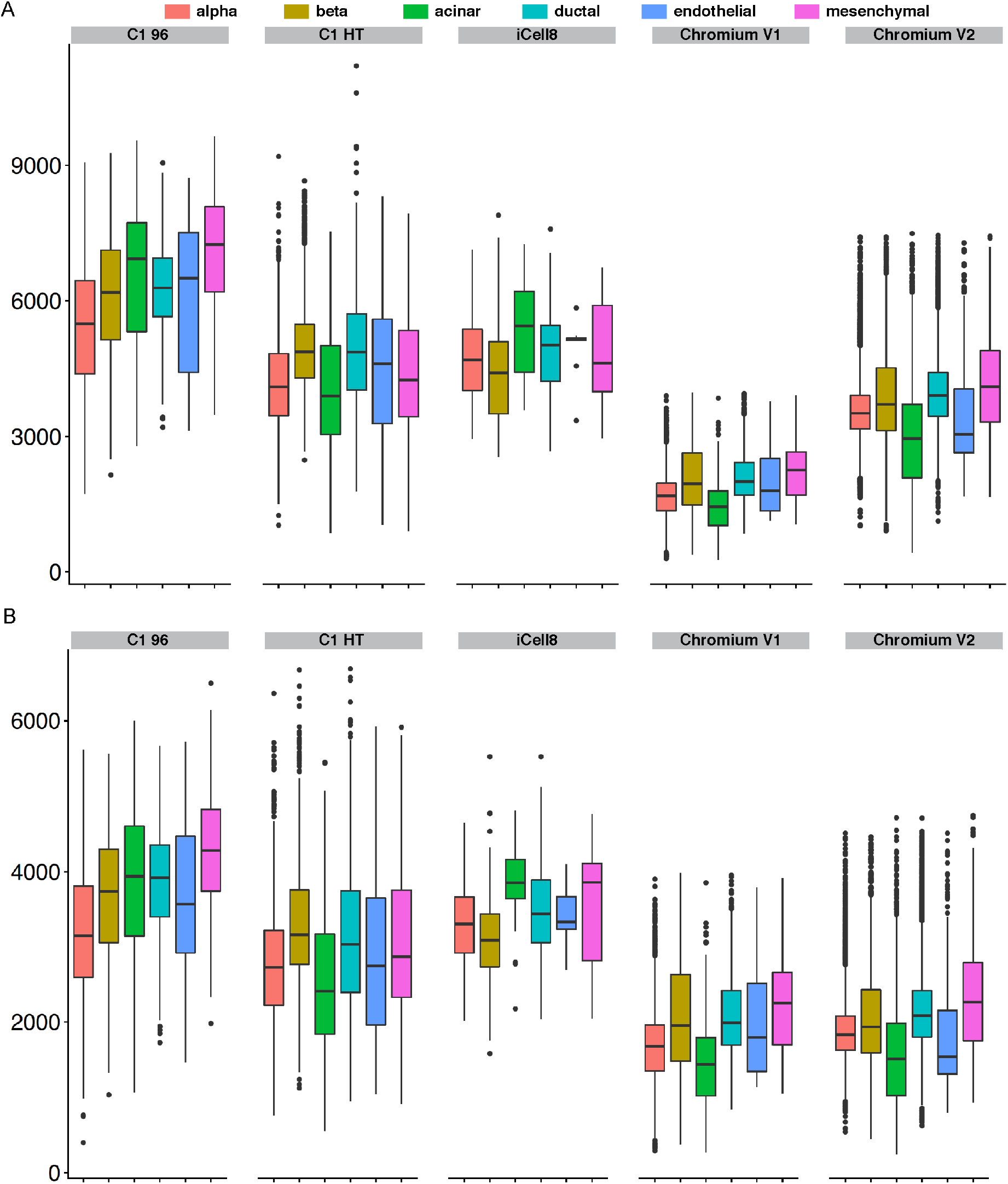
Sensitivity assessments. (A) Boxplots depict number of genes detected with at least 1 count or 1 UMI in different single-cell RNA-seq platforms for different pancreatic cell types. (B) Same as in (A), except that reads per cell from all platforms are down-sampled to comparable levels as in Chromium V1.

Next, we employed two orthogonal methods to evaluate the dropout frequency of the single-cell RNA-seq platforms. As expected, the dropout rate decreased with increased gene expression level (Figure S4). The relationship between dropout rate and gene expression level can be modeled with the Michaelis-Menten (MM) equation, where the MM constant (Km) reflects the efficiency of the underlying enzymatic reaction (Andrews and Hemberg, 2018). We observed the smallest Km in the C1 96 platform, followed by iCell8, Chromium V2 and C1 HT (Figure S4). The highest Km value was consistently observed in the Chromium V1 platform for all cell types assessed, indicating inefficiency in the reverse transcription reaction (Figure S4). In all platforms and for all cell types, the number of genes detected was reduced with increasing the threshold of the detection rate (Figure S5). To derive dropout rates, we calculated the proportion of cells with zero counts for those genes that were commonly expressed in at least 25% of the cells in at least one platform, as has been proposed before (Ziegenhain et al., 2017). Between 9,011 to 9,793 genes from each cell type were used for this calculation. Consistent with number of genes detected (Figure 4), the C1 96 system had the lowest dropout rate whereas Chromium V1 system had the highest (Figure 5A). Upon down-sampling of all single-cell reads to similar read depth as in Chromium V1, we again observed the lowest dropouts in C1 96, followed by iCell8 and C1 HT (Figure S6). Chromium V1 and V2 had comparable dropout rates with comparable sequencing depth (Figure S6). While the above method gives an overview of the relative dropout rate of the different single-cell platforms, it systematically underestimates the dropout rate since it does not account for those genes that were missed by all systems.

**Figure 5.**
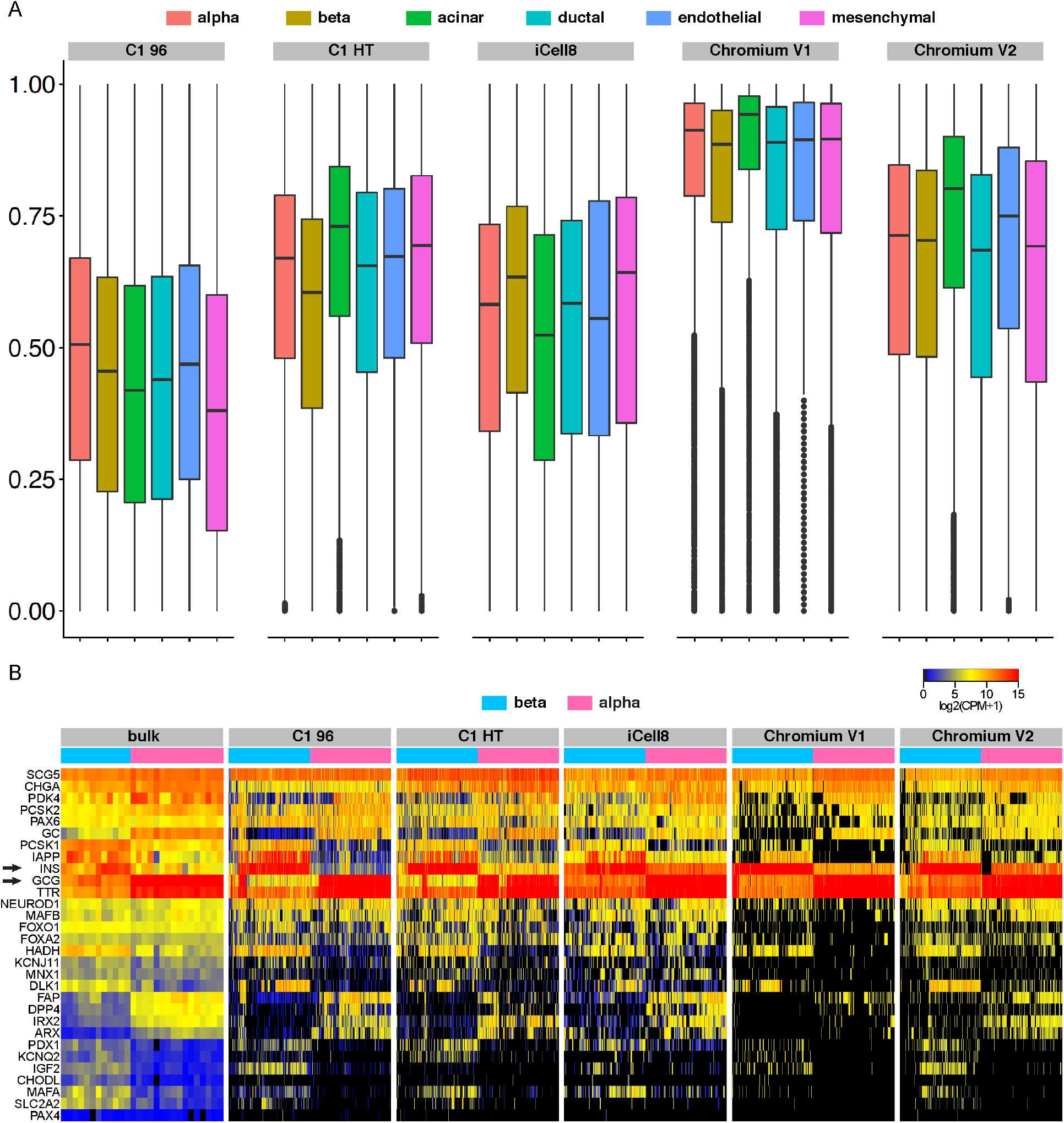
Dropout evaluations. See also Figures S4, S5 and S6. (A) Boxplots depict dropout rates in different single-cell RNA-seq platforms for different pancreatic cell types. Dropout rates are calculated based on proportions of cells with zero counts for genes that are commonly expressed in at least 25% of the cells in at least one platform. (B) Heatmaps display relative expression levels for selected markers in alpha and beta cells in different systems. Each column represents one sample (bulk) or one cell (all the singlecell platforms). Alpha and beta cells have variable but detectable levels of conflict expression of INS and GCG (arrows), correspondingly.

As a second way to evaluate dropout events, we employed a list of canonical markers for human alpha and beta cells to construct gene expression heatmaps (Figure 5B). As a reference, we included bulk RNA-seq data from sorted human alpha and beta cells (Ackermann et al., 2016). Compared with bulk (Figure 5B, left panel), all single cell platforms had frequent events of zero reads, as manifested by blocks of black color on the heatmap (Figure 5B). On the other hand, cell-type restricted marker genes displayed a more binary pattern in the single-cell platforms (Figure 5B). This phenomenon underscores the value of single-cell RNA-seq for obtaining purer gene expression signatures. One exception was the observed ‘conflicted’ expression of the signature hormones in alpha and beta cells (i.e. the detection of INS transcripts in alpha cells and of GCG transcripts in beta cells) (Figure 5B, black arrows). In both the C1 96 and C1 HT platforms, alpha and beta cells had lower conflicted hormone expression levels. However, alpha and beta cells in the iCell8, Chromium V1 and Chromium V2 platforms exhibited higher conflict hormone expressions compared with bulk. The platform specific discordant gene expressions indicated that this observation was likely to be technical rather than biological. We previously applied single-cell data from empty wells in the C1 96 platform to model the background contamination and concluded that the low level of conflicted hormone expression had likely originated from cell lysis events and the subsequent inclusion of ambient mRNA in all capture sites (Wang et al., 2016a). Similar processes likely also occur in the droplet-based systems.

### Donor batch effect

Due to the unpredictable nature of human donor availability, oftentimes we could not process multiple samples in one single-cell experiment. Therefore, we examined the potential donor and batch effects in our single-cell data. We selected alpha cells for this analysis since this cell type was the most abundant. We clustered alpha cells from each platform using graph-based approaches and visualized cluster identifiers together with donor labels in each platform using tSNE (Figure 6) (Butler et al., 2018). For the iCell8 and Chromium V1 systems, we only sequenced cells from one donor and therefore different clusters represent cellular heterogeneity rather than batch effects (Figure S7). For the remaining platforms, we observed various degrees of donor effects (Figure 6). Alpha cells analyzed by the C1 96 platform were separated into two clusters, and cells from different donors were scattered between these two clusters (Figure 6A).

**Figure 6.**
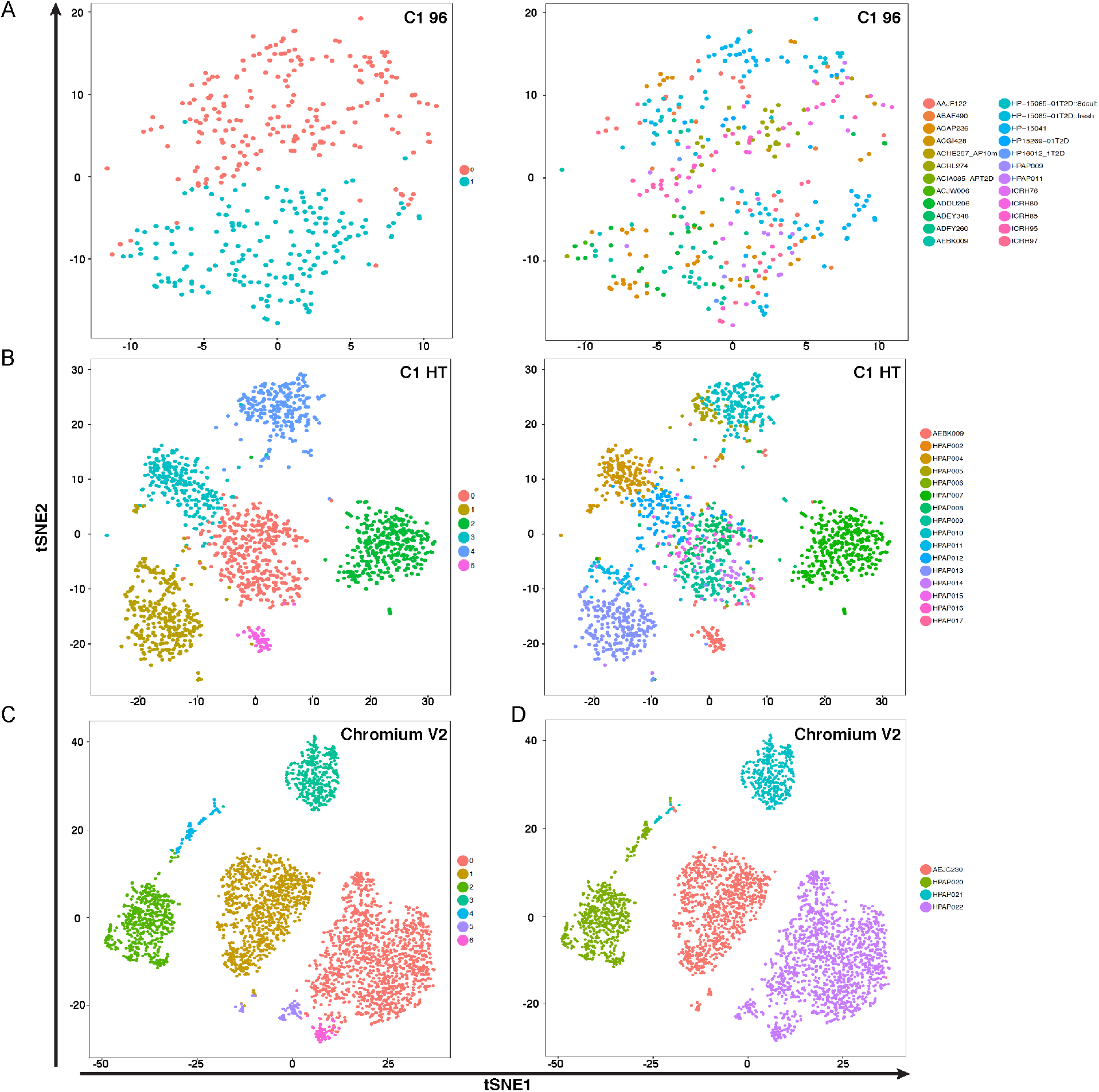
tSNE graphs for all the alpha cells in different single-cell platforms. See also Figure S7. Cluster labels are shown on the left-side tSNE plot in each panel. Donor labels are shown on the right-side tSNE plot in each panel. (A) Alpha cells from different donors distribute in two clusters in the C1 96 system. (B and C) Alpha cell clusters are driven by donors in the (B) C1 HT and (C) Chromium V2 systems.

On the other hand, in both the C1 HT and Chromium V2 systems, the alpha cell clusters were highly donor-specific (Figures 6B and 6C). These donor batch effects could potentially confound downstream analyses. One illustrative example is shown in Figure 3. When we performed cell type identification with all cells, because of the donor effects in the C1 HT and Chromium V2 platforms, alpha cells from different donors split into different clusters (Figures 3B and 3E). Without the guidance of marker gene expression, we might have erroneously concluded that these different clusters corresponded to different cell types.

### Differential expression analysis between alpha and beta cells from the same donor captured by different single-cell RNA-seq platforms

To further illustrate the impact of different single-cell technologies on biological discovery, we performed single-cell RNA-seq experiments on different systems using cells from a single donor, AEBK009. In particular, we utilized platforms spanning the spectrum of the commercially available single-cell systems: C1 96, with the highest sensitivity and the lowest throughput, C1 HT, with intermediate sensitivity and intermediate throughput, and Chromium V1, with the lowest sensitivity and highest throughput. As expected, the number of cells captured differed greatly by platform: in C1 96 platform, 27 alpha and 3 beta cells were obtained; in C1 HT platform, 61 alpha and 29 beta cells were acquired; and in Chromium V1 platform, 2,361 alpha and 974 beta cells were distinguished. Gene-level differential expression analyses between beta and alpha cells of individual platforms were carried out using the limma voom method, with a FDR cutoff of 10% (Law et al., 2014). With the increased cell number analyzed in the Chromium V1 system, more genes were identified as being differentially expressed (Figures. 7A and 7B). However, overall there was only a small number of concordantly differentially expressed genes identified by the three platforms (Figures 7A and 7B). We further crossreferenced our single-cell differential expression analyses with bulk RNA-seq data acquired from sorted human alpha and beta cells (Ackermann et al., 2016). Among the 22 genes identified in all three single-cell platforms as commonly upregulated in alpha cells, DPP4, FAP, GC, GCG, IGFBP2, MUC13, TM4SF4, TMEM45B and TTR were also identified by bulk RNA-seq analysis (Figure 7A, genes in red and Table 1 from Ackermann et al., 2016). Among the 8 genes commonly observed as upregulated in beta cells in the three single-cell platforms, BMP5 and INS were also enriched in the bulk assay (Figure 7B, genes in red and Table 1 from Ackermann et al., 2016). The large degree of discrepancy between the single-cell RNA-seq platforms, and between single-cell and bulk systems reflects the current technical limitations of the technology (see also Wang and Kaestner, 2019).

**Figure 7.**
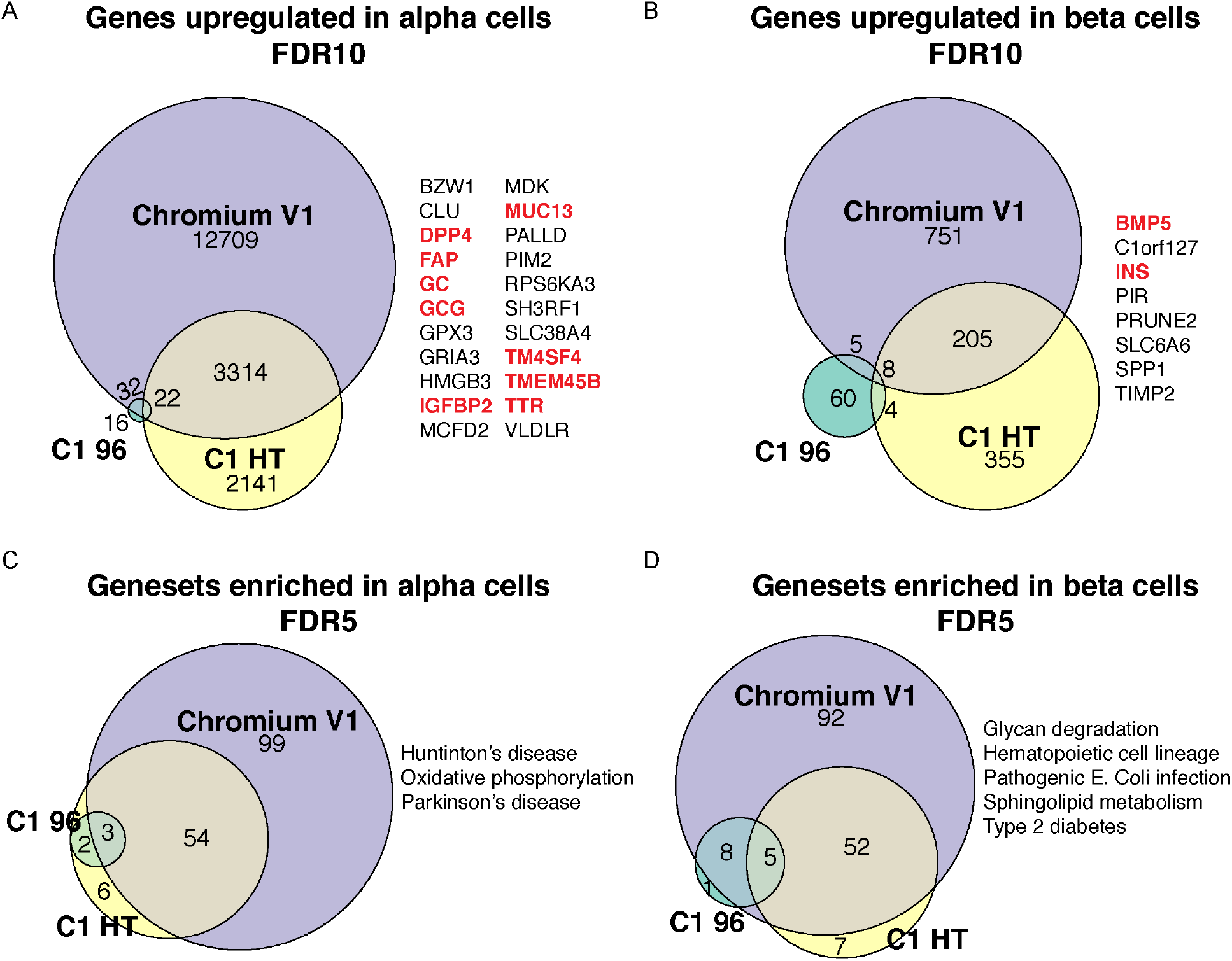
Comparative analyses of alpha and beta cells isolated from one donor in three different single-cell RNA-seq platforms. (A-B) Venn diagrams summarize results from gene level differential expression analyses. FDR<0.1 is used as statistical cutoff to select genes that are upregulated in (A) alpha or (B) beta cells. Genes that are consistently detected as upregulated in all three platforms are listed on the right. Red font indicates concordant genes identified also in bulk data (Ackermann et al., 2016). (C-D) Venn diagrams summarize results from geneset level enrichment analyses. FDR<0.05 is used as statistical cutoff to select gene sets that are upregulated in (C) alpha or (D) beta cells. Gene sets that are consistently detected as upregulated in all three platforms are listed on the right.

Next, we performed gene set enrichment analyses using Gene Set Variation Analysis (GSVA) aiming to buffer some of the noises from single gene analyses (Hanzelmann et al., 2013). Differentially expressed genes were used to compute the GSVA scores for 186 KEGG canonical pathway gene sets (Molecular Signatures Database v6.2) (Subramanian et al., 2005). We observed that the congruent information among different single-cell platforms increased when comparing gene sets (Figures 7C and 7D). Huntinton’s disease, Oxidative phosphorylation and Parkinson’s disease pathways were the three gene sets that were identified by all three platforms to be enriched in alpha cells (Figure 7C). Glycan degradation, Hematopoietic cell lineage, Pathogenic *E. Coli* infection, Sphingolipid metabolism and Type 2 diabetes were among the five gene sets that were consistently identified to be enriched in beta cells (Figure 7D).

### Cost estimate and efficiency comparison

Based on actual expenses incurred, we calculated the average cost of single-cell experiments for different platforms (Table 2). Our calculation excluded costs of obtaining islets, isolating single cells, consumable plastics as well as labor costs, and costs of all reagents were based on manufacturers’ catalogs. Per experiment, the expenses for single-cell library generation and sequencing for C1 96 platform were the lowest (~$3,500). C1 HT, Chromium V1 and Chromium V2 platforms had similar costs per experiment (~$5,600). The iCell8 platform had the highest cost (~$7,600), mostly due to the expense of the non-reusable alloy wafer. On the other hand, at a per cell level, C1 96 platform was the most expensive one (~$83/cell). C1 HT and iCell8 system had comparable price (~$17/cell). And Chromium V1 and Chromium V2 platforms had the lowest expenses (~$1/cell). To be noted, our cost per cell estimate was based on cost per ‘good’ cell (cells passed QC). As a result, our price estimates were higher than what is reported by the manufacturers, whose estimates are based on ideal scenarios with 100% efficiency.

**Table 2.**
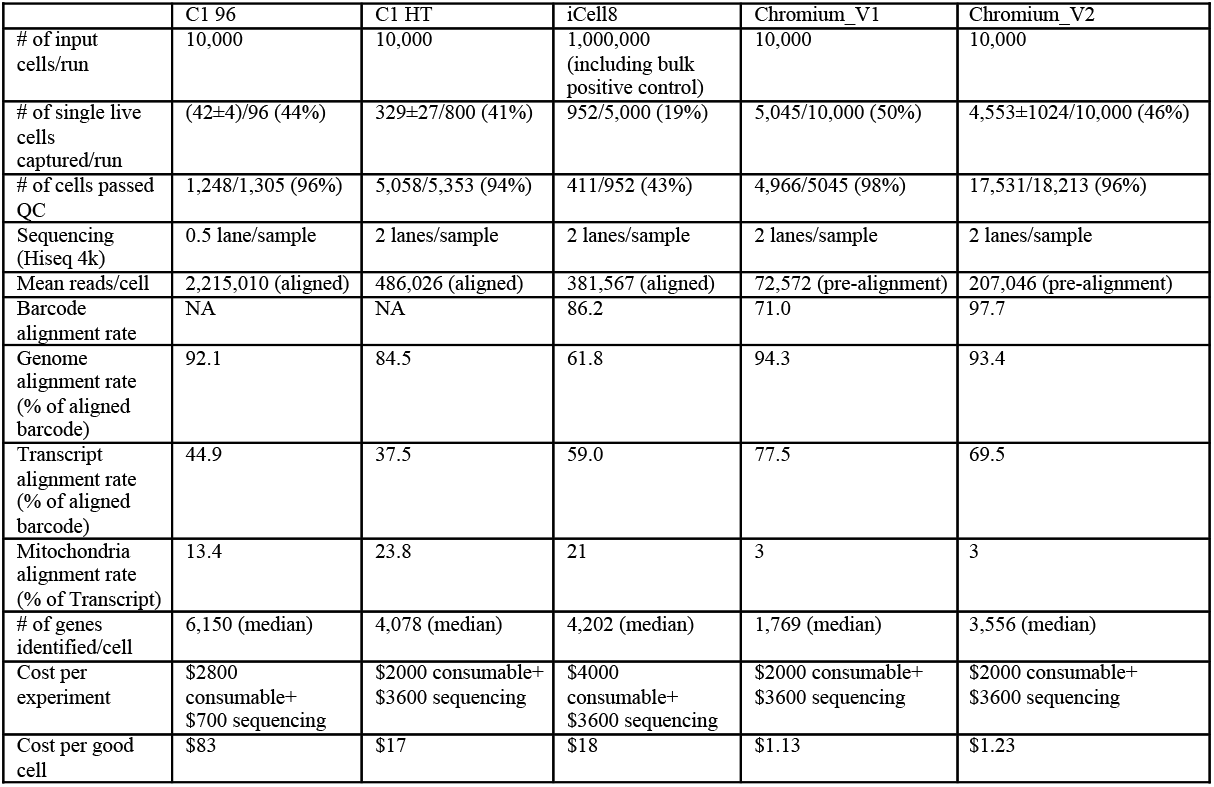
Summary of single-cell RNA-seq platform statistics. Related to Figure 1.

|  | C1 96 | C1 HT | iCell8 | Chromium V1 | Chromium V2 |
| --- | --- | --- | --- | --- | --- |
| # of input cells/run | 10,000 | 10,000 | 1,000,000 (including bulk positive control) | 10,000 | 10,000 |
| # of single live cells captured/run | (42±4)/96 (44%) | 329±27/800 (41%) | 952/5,000 (19%) | 5,045/10,000 (50%) | 4,553±1024/10,000 (46%) |
| # of cells passed QC | 1,248/1,305 (96%) | 5,058/5,353 (94%) | 411/952 (43%) | 4,966/5045 (98%) | 17,531/18,213 (96%) |
| Sequencing (HiSeq 4k) | 0.5 lane/sample | 2 lanes/sample | 2 lanes/sample | 2 lanes/sample | 2 lanes/sample |
| Mean reads/cell | 2,215,010 (aligned) | 486,026 (aligned) | 381,567 (aligned) | 72,572 (pre-alignment) | 207,046 (pre-alignment) |
| Barcode alignment rate | NA | NA | 86.2 | 71.0 | 97.7 |
| Genome alignment rate (% of aligned barcode) | 92.1 | 84.5 | 61.8 | 94.3 | 93.4 |
| Transcript alignment rate (% of aligned barcode) | 44.9 | 37.5 | 59.0 | 77.5 | 69.5 |
| Mitochondria alignment rate (% of Transcript) | 13.4 | 23.8 | 21 | 3 | 3 |
| # of genes identified/cell | 6,150 (median) | 4,078 (median) | 4,202 (median) | 1,769 (median) | 3,556 (median) |
| Cost per experiment | \$2800 consumable+ \$700 sequencing | \$2000 consumable+ \$3600 sequencing | \$4000 consumable+ \$3600 sequencing | \$2000 consumable+ \$3600 sequencing | \$2000 consumable+ \$3600 sequencing |
| Cost per good cell | \$83 | \$17 | \$18 | \$1.13 | \$1.23 |

## DISCUSSION

In this study, we systematically compared three major commercial single-cell RNA-seq platforms using data generated from primary human islets. The human islets represent a “real-world” example from which the performance of different single-cell platforms could be directly translated to other complex biological systems and primary cells.

In human islet samples, we observed higher doublets rates (21-33%, Figure 2) than typically reported using mixed species experiments. Since we used proportions of INS and GCG double positive cells to estimate the doublet rates, these numbers likely represent an overestimation of the true doublet rate due to the facts that: (1) some INS+/GCG+ doublets are naturally occurring within human islets; (2) INS+ (beta) and GCG+ (alpha) cells are frequently directly juxtaposed to each other in the pancreatic tissue. As such, they are prone to be doublets resulting from incomplete dissociation of cells from the primary tissue source; and (3) human pancreatic endocrine cells are epithelial cells with a high propensity to re-aggregate in solution. Thus, caution should be taken to ensure good single-cell isolation and to filter out doublets events when performing single-cell experiment on cells from primary tissues.

In our analyses, reaction chamber-based technologies (C1 96, C1 HT, and iCell8) displayed higher sensitivity and lower dropout rates compared with droplet-based technologies (Chromium V1 and Chromium V2) (Figures 4A and 5). Among the reaction chamber-based technologies, C1 96 platform has the best performance over a variety of cell types (including epithelial, endothelial and mesenchymal cells) (Figures 4A, 5, and S4–S6). Part of the increase in detection efficiency can be explained by the relatively higher sequencing depth in the chamber-based technologies (Table 2). However, even after down-sampling of all the single-cell data to similar read depth as achieved by the Chromium V1 system, reaction chamber-based platforms still exhibited higher sensitivity and lower dropout frequency (Figures 4B and S6). This observation suggests that reaction chamber-based platforms have higher efficiency in capturing cellular mRNA molecules. It is noteworthy that different cell types displayed varying levels of detectable RNA transcripts, and that the relative gene counts were not consistent among different single-cell RNA-seq platforms (Figure 4). For example, acinar cells are among the cell types with the highest numbers of genes detected in the C1 96 and iCell8 platforms but are among the cell types with the lowest numbers of genes detected in the C1 HT, Chromium V1 and Chromium V2 platforms (Figure 4). The difference in gene diversity by cell type observed in different platforms may indicate underlying amplification bias in different platforms.

Among the three single-cell platforms where we processed multiple donors, cells clustered based on their donors of origin in the C1 HT and Chromium V2 platforms; whereas in the C1 96 platforms, cells from different donors are homogeneously distributed in multiple clusters (Figure 6). The large batch effects observed in C1 HT and Chromium V2 platforms result from a combination of donor and technical variation. The reduced level of batch effect associated with the C1 96 platform suggests that a more sensitive platform with deeper sequencing depth can mitigate the variation to a large degree. Recently, several algorithms were developed specifically to deal with single-cell batch effects and to align different datasets (Butler et al., 2018; Haghverdi et al., 2018).

The minimal congruency between different single-cell platforms when we compared alpha and beta cells from the same donor highlights the technical variability in single-cell RNA-seq experiments (Figure 7). High proportions of dropouts in the single-cell data contribute to this variation (Figure 5) (Hicks et al., 2018). This high technical variability among different singlecell platforms, together with the high variability among different samples within the same singlecell platform, presents a serious challenge to the interpretation of single-cell RNA-seq data. Future technical and computational advancements, combined with larger single-cell RNA-seq datasets, promise to reduce the technical artefacts and allow for the identification of true biological phenotypes (Wang and Kaestner, 2018).

In summary, we performed comprehensive analyses of three major commercial singlecell RNA-seq platforms using primary human islet cells. We showed that the C1 96 platform has the highest sensitivity, lowest throughput and the highest experimental expense per cell among all platforms. On the other end of the spectrum, the Chromium system has the lowest sensitivity, highest throughput and the lowest experimental cost per cell. We demonstrated noticeable doublets rates, large donor batch effects and high technical variation among different single-cell RNA-seq platforms. Our analyses provide a reference for future single-cell RNA-seq studies, and indicate that single cell RNA-seq technology has not yet reached full maturity.

## MATERIAL AND METHODS

### Human pancreatic islet sample preparation

Human islets were obtained through the Diabetes Research Center of the University of Pennsylvania (NIH DK 19525), the Integrated Islet Distribution Program (IIDP) (http://iidp.coh.org/) and the HPAP consortium (https://hpap.pmacs.upenn.edu/) of the Human Islet Research Network (https://hirnetwork.org/). 200 to 500 hand-picked islets were digested in 0. 05%Trypsin-EDTA (ThermoFisher 25300054) for up to 9 min at 37°C, with physical dispersion by p1000 pipette every 2 min. Islets dissociation was monitored under a microscope and was stopped by PBS +10% FBS when 90% of the cells were released into single cells. In the C1 96, C1 HT and iCell8 system, dissociated pancreatic cells were further labeled with viability and/or nuclear dye (see Single-cell RNA-seq library preparation below). Cells were counted with a Countess Automated Cell Counter (ThermoFisher C10227) and diluted to appropriate concentrations for each platform (see Single-cell RNA-seq library preparation below). Immediately before loading, cells were filtered through a 35 μm cell strainer (Corning 352235) to remove clumps.

### Single-cell RNA-seq library preparation

#### Fluidigm C1

Single cells were stained with 4 μM Ethidium homodimer-1 and 2 μM Calcium AM (ThermoFisher LIVE/DEAD Viability Kit L3224) in the dark at room temperature for 20 min. Stained cells were washed once with PBS + 10% FBS and 10 μl of the cell suspension was loaded onto a C1 96-IFC (Fluidigm 100-5760) or C1 800HT-IFC (Fluidigm 101-4982) at a concentration of 1,000 cells/μl. Single-cell RNA-seq libraries were prepared according to the manufacturer’s protocol (Fluidigm PN100-7168 K1 and PN101-4964 B1). 10-17 μm IFCs were used. Cell loading was scanned by Keyence BZ-x710 at 10x magnification using the channels of GFP, TRITC and bright field. Single live cells were curated manually.

#### Clontech iCell8

Single cells were stained with Hoechst 33342 and Propidium Iodide according to the manufacturer’s protocol (Clontech iCell8 D07-000040-004 rev 4). Before cell loading, 50,000 cells were separated and RNA was extracted from this population using Trizol. RNA from this preparation served as the bulk positive control. 100 μl of cells at a concentration of 20 cells/μl was loaded in 8 wells (A1 to D2) of a 384-well plate and dispensed into the iCell8 chip. The chip was scanned and analyzed using the iCell8 build-in microscope and CellSelect software. 952 wells automatically annotated to contain single live cells, together with 88 additional wells labeled as “double”, “dead”, “small”, “large”, “multiple”, or “no” cells were manually selected for subsequent library preparation and sequencing. In addition, 5 positive (RNA from parallel processed bulk sample) and 5 negative (PBS) on-chip controls were included for downstream library preparation and analyses.

#### 10x Genomics Chromium

10,000 single cells were loaded in the 10x Single Cell 3’ Chip at a concentration of 1,000 cells/ μl. For the V1 chemistry, library preparation was carried out according to 10x Genomics CG00026 RevB. For the V2 chemistry, library preparation was performed according to 10x Genomics CG00052 RevA.

### Sequencing geometry and reads alignment

The C1 96 libraries were sequenced on the Illumina HiSeq 2500 system with 100-bp single-end reads. The Chromium V1 library was sequenced on a combination of HiSeq 2500 with rapid mode and NextSeq sequencer with 98×26 paired-end reads. All the remaining single-cell libraries were sequenced on the Illumina HiSeq 4000 system. The C1 800HT libraries were sequenced with 25×75 paired-end reads. The iCell8 library was sequenced with 25×100 paired-end reads. The Chromium V2 libraries were sequenced with 26×98 paired-end reads.

Sequencing data from single-cell libraries were processed using optimal aligners for each platform. For the sequencing data from C1 96, read alignment and gene expression quantification were performed using RNA-seq unified mapper (RUM) (Grant et al. 2011). For C1 800HT data, mapping and quantification were performed by STAR (Dobin et al. 2013). For the iCell8 sequencing data, RSEM was used to derive transcript count (Li and Dewey 2011). The Chromium libraries were processed with Cell Ranger 2.2. The Fluidigm and iCell8 protocol did not include unique molecular identifiers (UMI). The 10x libraries had 3’UMI on the bead-bound oligo dT primers to facilitate accurate transcript counting.

### Cell type annotation and donor batch effect assessment

To annotate cell types, we implemented the Seurat R package (Butler et al., 2018) and selected the top variable genes for dimension reduction, followed by graphic based clustering and tSNE visualization. Cluster annotation was based on canonical cell type markers, including GCG for alpha cells, INS for beta cells, SST for delta cells, KRT19 for ductal cells, PRSS1 for acinar cells, PECAM1 for endothelial cells, and SPARC for mesenchymal cells. In the iCell8 system, basic Seurat process did not resolve cell type clusters. Thus, consensus clustering SC3 method with k=8 was used to identify cells of different cell types (Kiselev et al., 2017). The cell type labels from SC3 were projected to the Seurat tSNE graph for visualization. For donor batch effect assessment, alpha cells from all single-cell platforms were extracted. Graphic based clustering and tSNE visualization were performed similarly as in cell type annotation process outlined above.

### Sensitivity and dropout rates estimation

The number of genes detected with at least one read or one UMI was used to measure sensitivity. The Michaelis-Menten constant (Km) for Dropout versus Expression level relationship was estimated with the M3Drop package (Andrews and Hemberg, 2018). To derive dropout rates, the proportion of cells with zero counts for those genes that were commonly expressed in at least 25% of the cells in at least one platform was calculated. The heatmap with canonical alpha and beta cell maker genes was constructed using heatmap.2 function in gplots package, with “ward.D2” for hclustfun.

### Differential expression analyses

Gene level differential expression analyses were performed with the limma voom method (Law et al., 2014). 186 genesets from c2.cp.kegg.v6.2.symbols.gmt were used to compute GSVA scores for geneset level enrichment analyses (Hanzelmann et al., 2013; Subramanian et al., 2005). Statistical significance was subsequently computed by the limma package (Ritchie et al., 2015).

## AUTHOR CONTRIBUTIONS

Y.J.W. acquired, researched the data and wrote the manuscript. J.S., J.L., Z.W., and A.K. performed sequencing data processing and analyses. The HPAP Consortium isolated human pancreata. K.H.K. conceived of the study, reviewed and edited the manuscript.

## ACKNOWLEDGEMENTS

The authors thank all the islet donors and their families who make this work possible. The authors also thank the University of Pennsylvania Diabetes Research Center for the use of the Functional Genomics (National Institute of Diabetes and Digestive and Kidney Diseases [NIDDK] P30-DK19525). This research was performed using resources and/or funding provided by the NIDDK-supported Human Islet Research Network (HIRN, RRID:SCR_014393; https://hirnetwork.org) grant UC4 DK104119 to K.H.K.

## Supplemental Information

**Figure S1.**
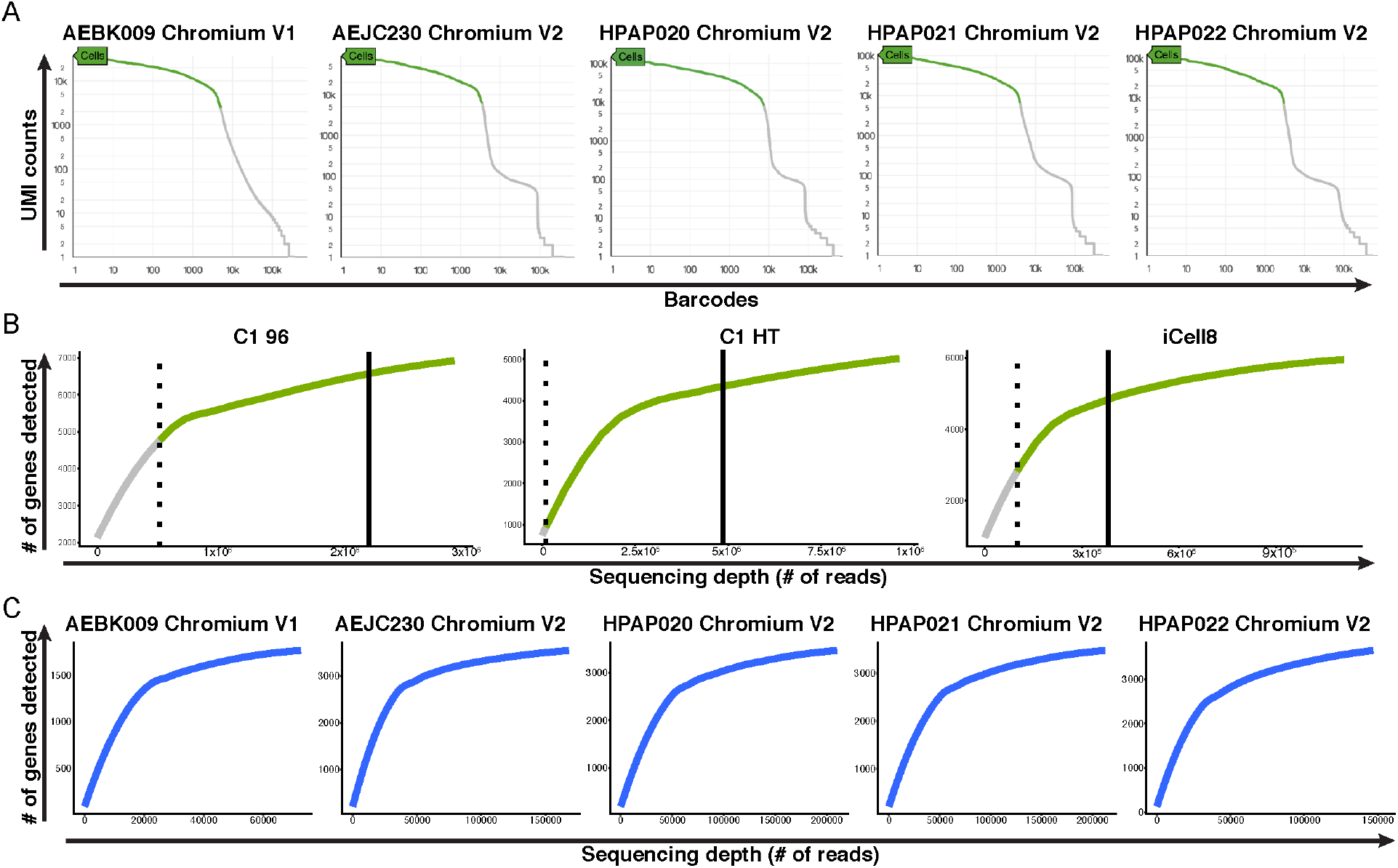
Quality control and data preprocessing. Related to Figure 1. (A) Cell filtering in the Chromium platform. Cells passing filters are represented on the green regions on each curve. (B) Loess regression curves display the relationships of number of genes detected with different sequencing depth in the C1 96, C1 HT and iCell8 systems. Single-cell data from all experiments on each system are combined. Within each graph, solid line indicates population median and dashed line indicates two standard deviation below population median. Cells passing filters are represented on the green regions on each curve. (C) Relationships of number of genes detected and varying sequencing depth in the Chromium system. Each curve is associated with one Chromium experiment. The endpoint of each curve represents the average sequencing depth for cells passing filter.

**Figure S2.**
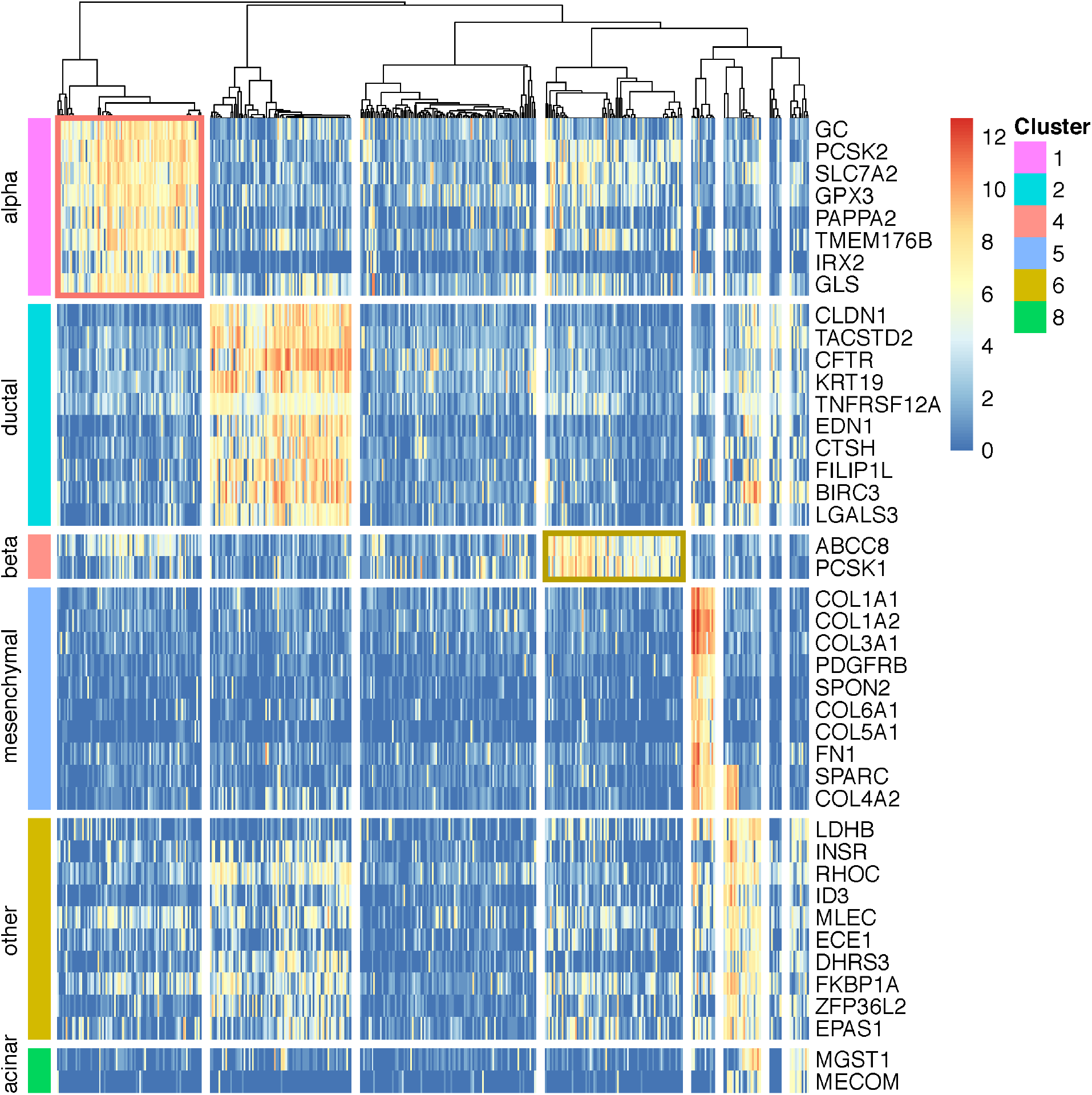
Heatmap displays relative levels of marker genes identified by SC3. Related to Figure 3. Six clusters are identified with SC3 algorithm (k=9). Two clusters are associated with alpha and beta cells (outlined by coral and gold colored rectangles respectively).

**Figure S3.**
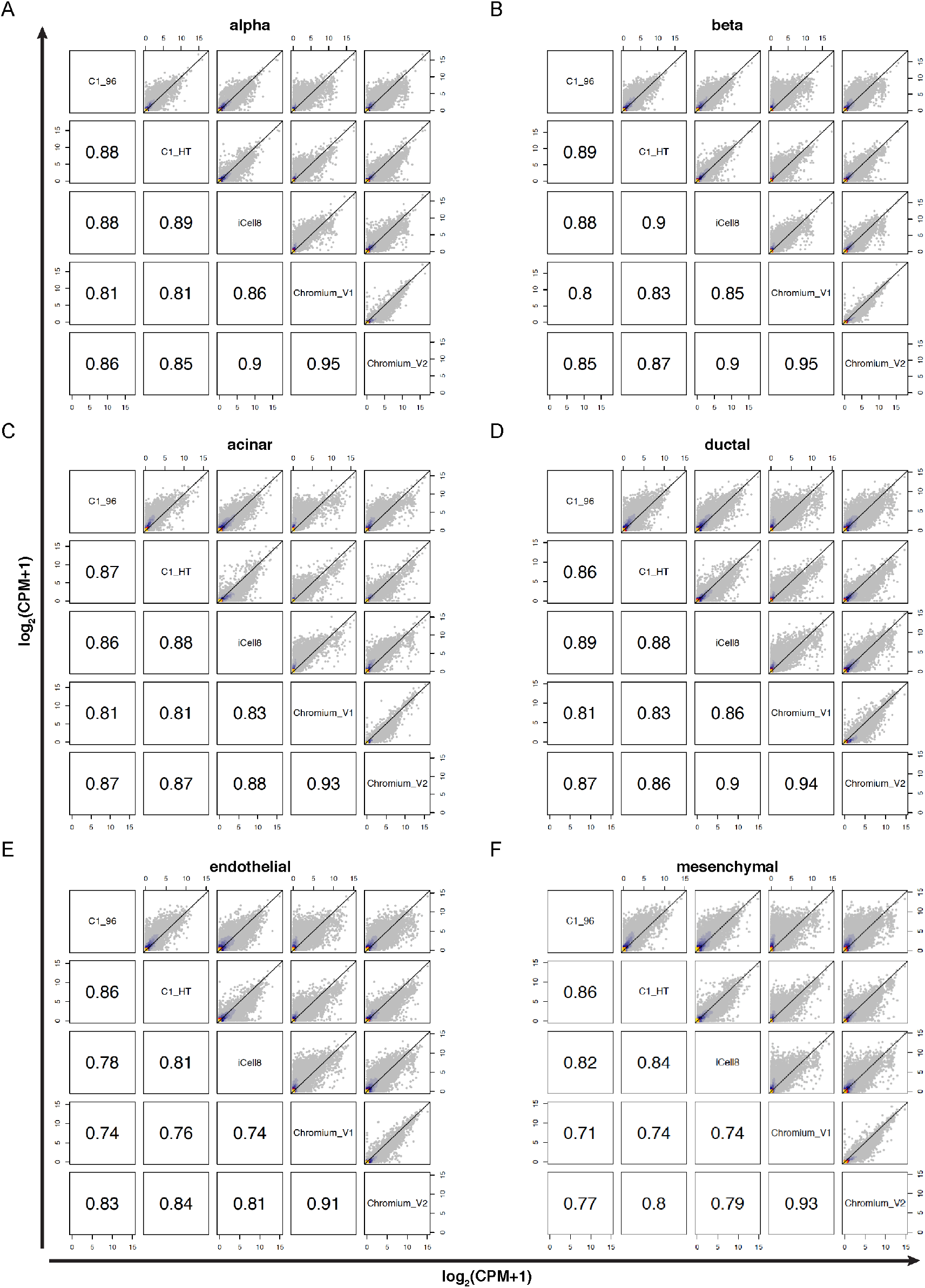
Pairwise correlation plots between all single-cell platforms in each cell type. Related to Figure 3. log2(CPM+1) for each gene is used to construct the scatter plots. Scatter plots correspond to (A) alpha, (B) beta, (C) acinar, (D) ductal, (E) endothelial, and (F) mesenchymal cells. Values shown in the lower triangle of each plot are spearman correlation between different pairs. Color in each scatter plot represents Kernel Density Estimation.

**Figure S4.**
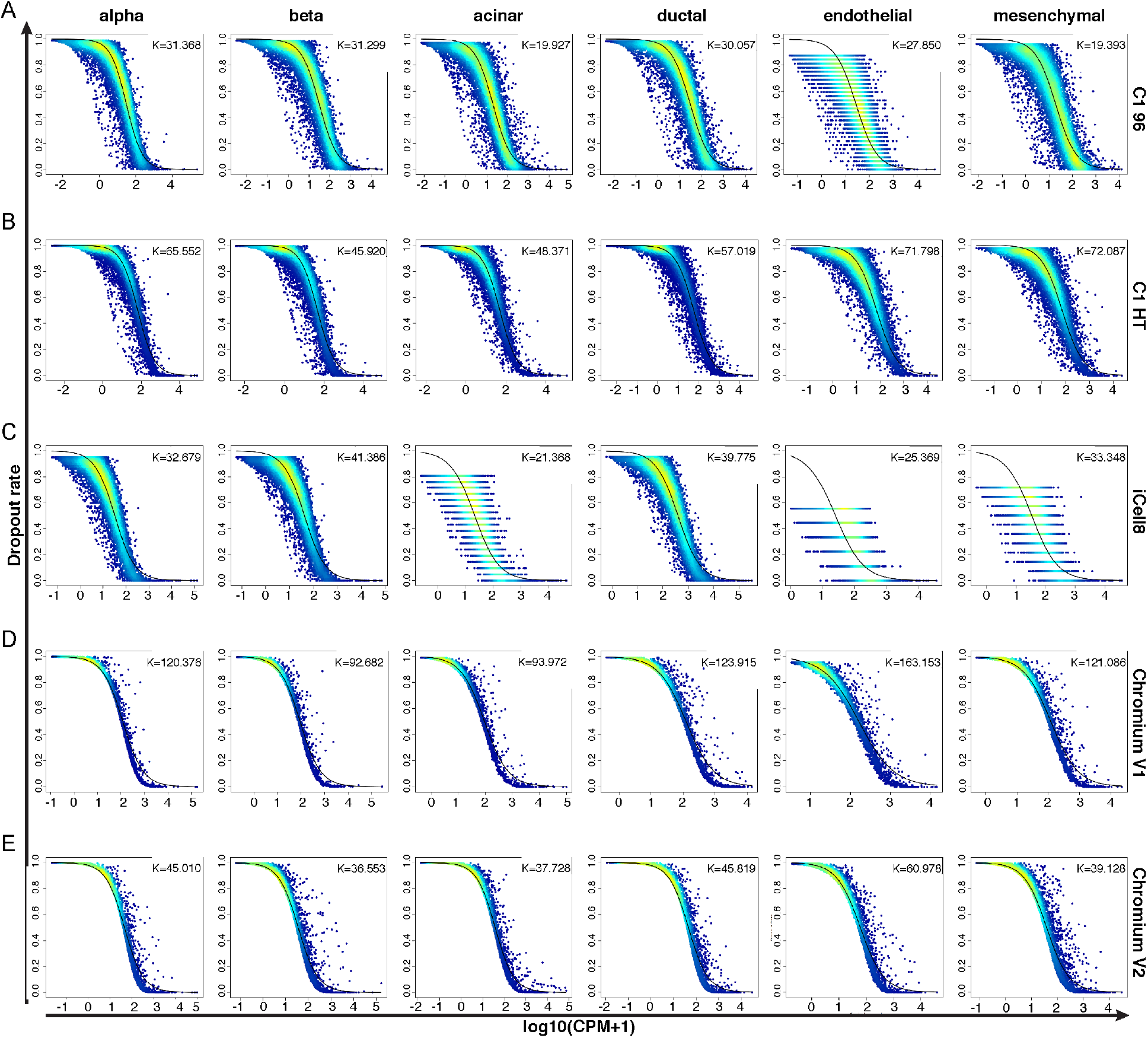
Michaelis-Menten (MM) curve illustrates the relationship between expression levels (log10(CPM+1)) and dropout rates. Related to Figure 5. MM plots for each cell type are shown for (A) C1 96, (B) C1 HT, (C) iCell8, (D) Chromium V1, and (E) Chromium V2 platforms. The MM constant (Km) for each fitting is shown on the upper right corner of each plot.

**Figure S5.**
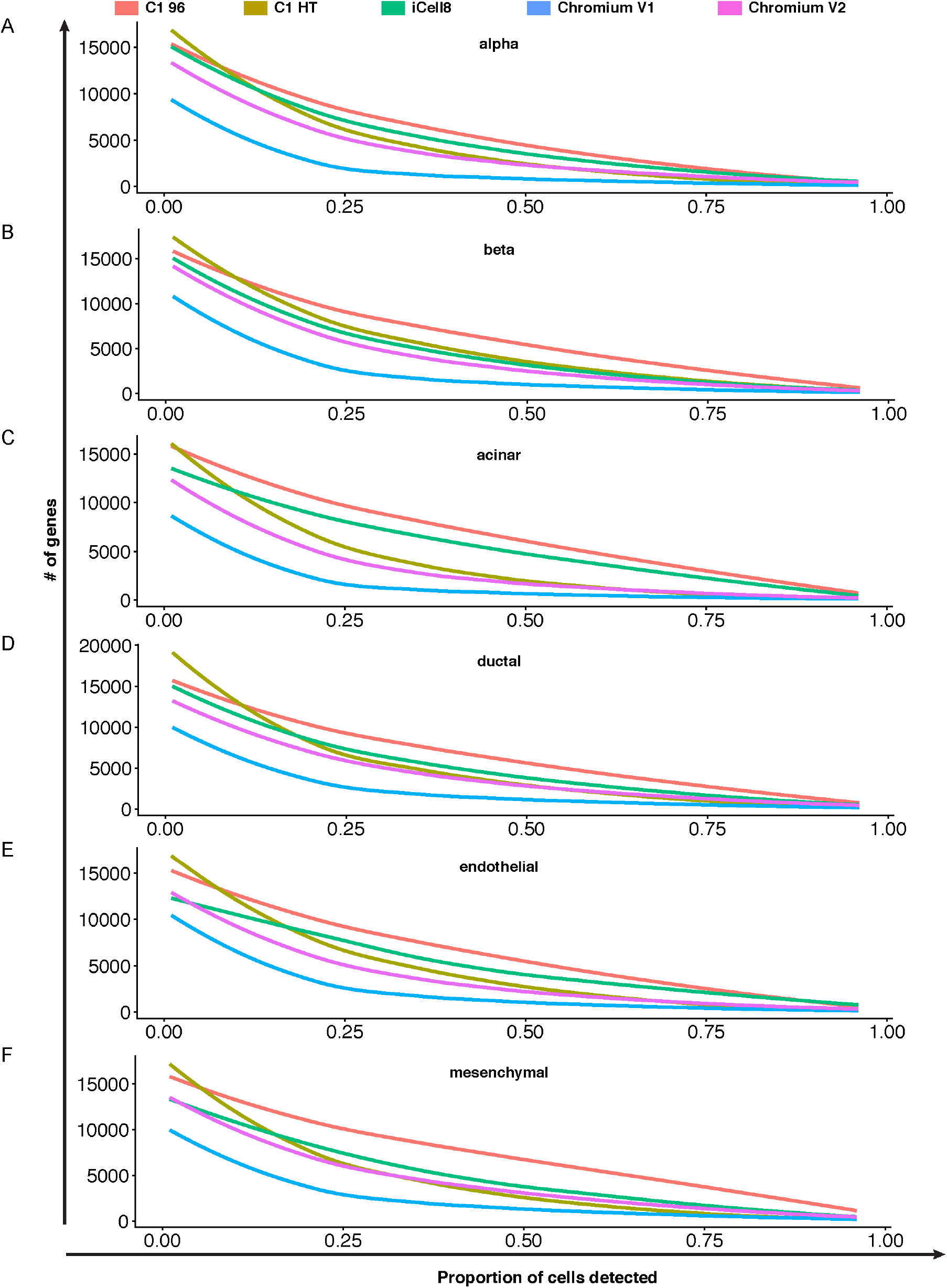
Loess curves display the dependency of number of genes detected on the threshold of the detection rate. Related to Figure 5. Regression curves are shown for (A) alpha, (B) beta, (C) acinar, (D) ductal, (E) endothelial, and (F) mesenchymal cells.

**Figure S6.**
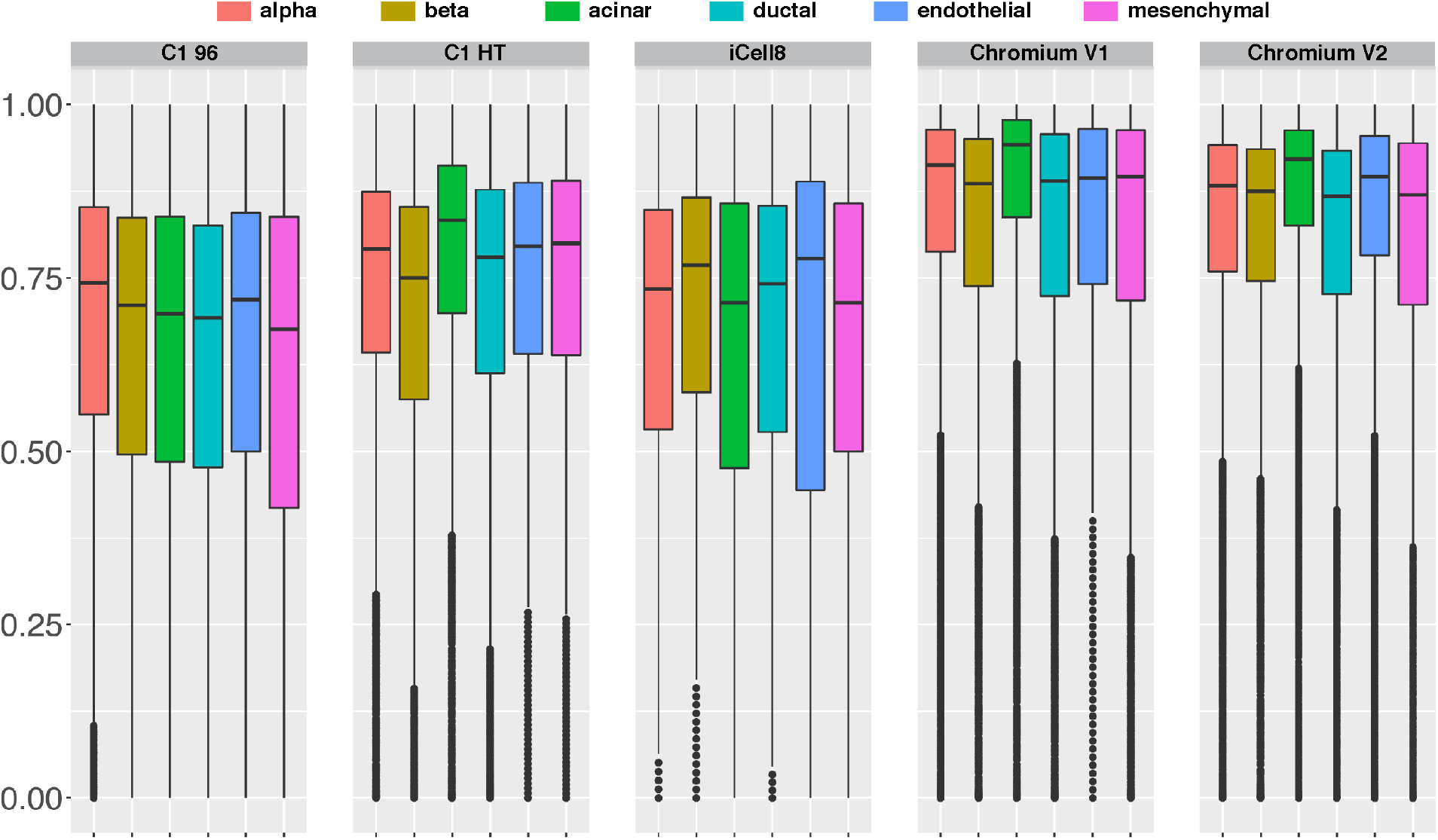
Dropout rate with down-sampling. Related to Figure 5. Reads from different platforms are down-sampled to be comparable levels as Chromium V1.

**Figure S7.**
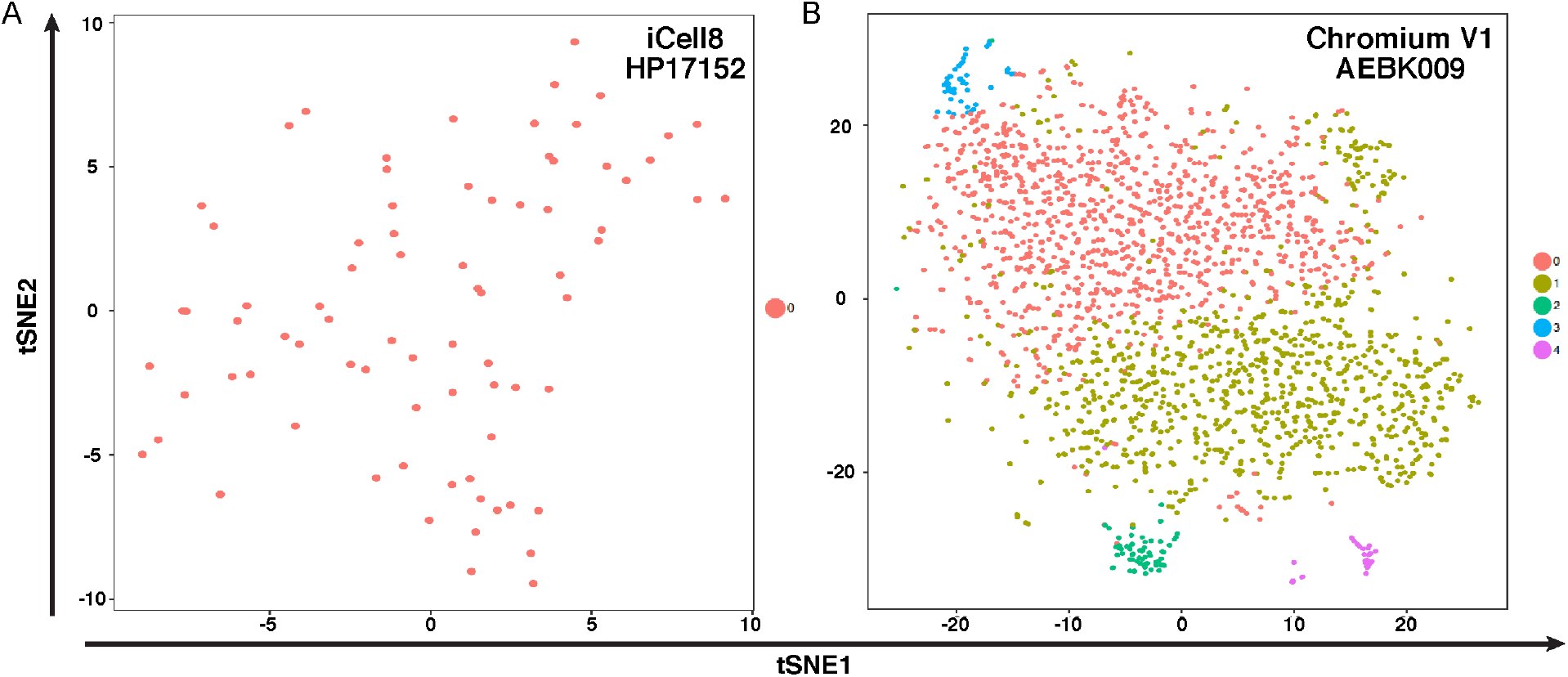
tSNE graphs demonstrate alpha cell heterogeneity in two of the single-cell platforms. Related to Figure 6. (A) Alpha cells identified in the iCell8 platform do not show distinct clusters. (B) Alpha cells from Chromium V1 platform show five distinct clusters.

**Table S1.**
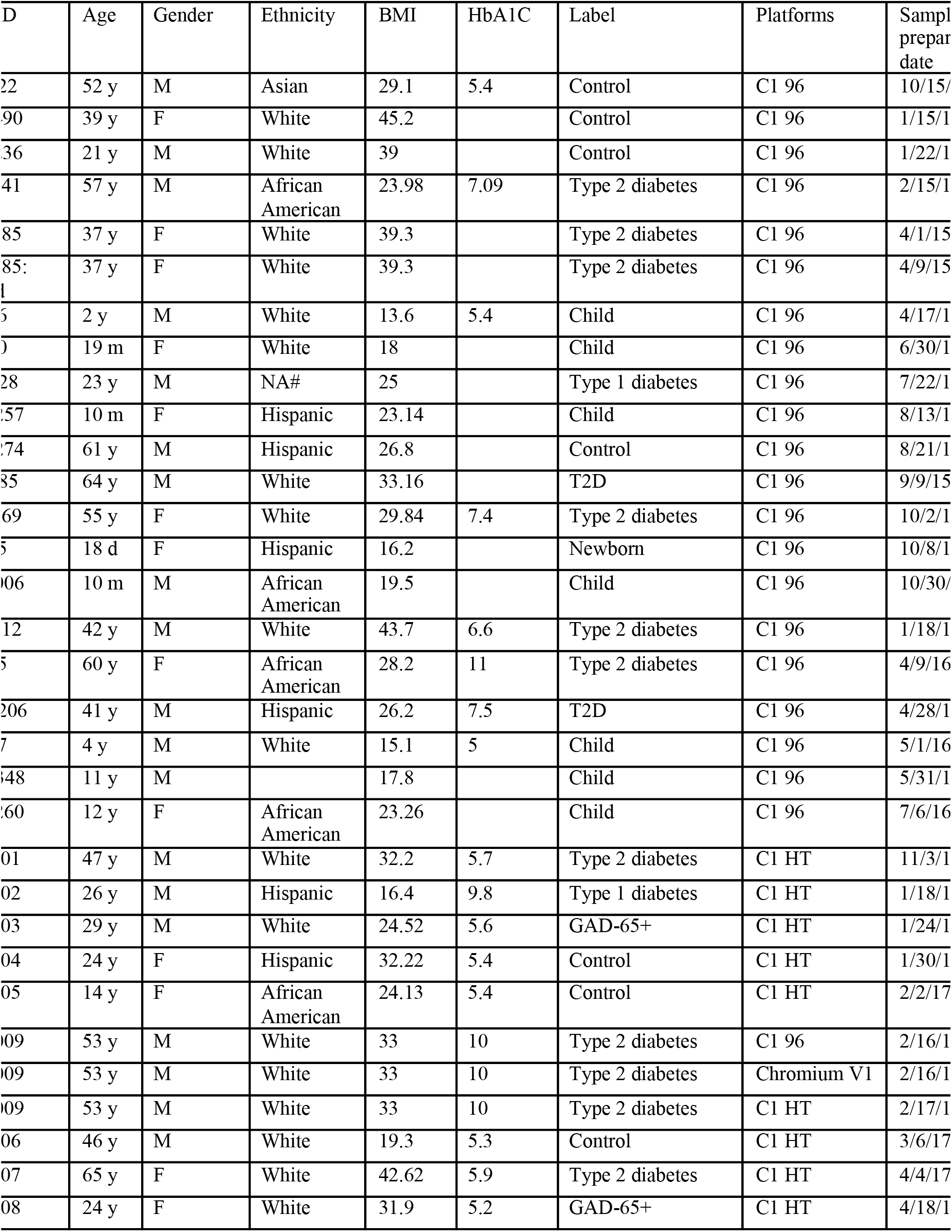
Donor information. Related to Figure 1.

## REFERENCES

1. Ackermann, A.M., Wang, Z., Schug, J., Naji, A., and Kaestner, K.H. (2016). Integration of ATAC-seq and RNA-seq identifies human alpha cell and beta cell signature genes. Mol Metab 5, 233–244.

2. Andrews, T.S., and Hemberg, M. (2018). M3Drop: Dropout-based feature selection for scRNASeq. Bioinformatics.

3. Butler, A., Hoffman, P., Smibert, P., Papalexi, E., and Satija, R. (2018). Integrating single-cell transcriptomic data across different conditions, technologies, and species. Nat Biotechnol 36, 411–420.

4. Dobin, Alexander, Carrie A. Davis, Felix Schlesinger, Jorg Drenkow, Chris Zaleski, Sonali Jha, Philippe Batut, Mark Chaisson, and Thomas R. Gingeras. 2013. “STAR: Ultrafast Universal RNA-Seq Aligner.” Bioinformatics 29 (1): 15–21.

5. Fluidigm Redesign of C1 Medium-Cell 96 and HT IFCs Improves Single-Cell Capture Efficiency.

6. Goldstein, L.D., Chen, Y.J., Dunne, J., Mir, A., Hubschle, H., Guillory, J., Yuan, W., Zhang, J., Stinson, J., Jaiswal, B., et al. (2017). Massively parallel nanowell-based singlecell gene expression profiling. BMC Genomics 18, 519.

7. Grant, Gregory R., Michael H. Farkas, Angel D. Pizarro, Nicholas F. Lahens, Jonathan Schug, Brian P. Brunk, Christian J. Stoeckert, John B. Hogenesch, and Eric A. Pierce. 2011. “Comparative Analysis of RNA-Seq Alignment Algorithms and the RNA-Seq Unified Mapper (RUM).” Bioinformatics 27 (18): 2518–28.

8. Haghverdi, L., Lun, A.T.L., Morgan, M.D., and Marioni, J.C. (2018). Batch effects in single-cell RNA-sequencing data are corrected by matching mutual nearest neighbors. Nat Biotechnol 36, 421–427.

9. Hanzelmann, S., Castelo, R., and Guinney, J. (2013). GSVA: gene set variation analysis for microarray and RNA-seq data. BMC Bioinformatics 14, 7.

10. Hicks, S.C., Townes, F.W., Teng, M., and Irizarry, R.A. (2018). Missing data and technical variability in single-cell RNA-sequencing experiments. Biostatistics 19, 562–578.

11. Ilicic, Tomislav, Jong Kyoung Kim, Aleksandra A. Kolodziejczyk, Frederik Otzen Bagger, Davis James McCarthy, John C. Marioni, and Sarah A. Teichmann. 2016. “Classification of Low Quality Cells from Single-Cell RNA-Seq Data.” Genome Biology 17 (February): 29.

12. Islam, S., Zeisel, A., Joost, S., La Manno, G., Zajac, P., Kasper, M., Lonnerberg, P., and Linnarsson, S. (2014). Quantitative single-cell RNA-seq with unique molecular identifiers. Nat Methods 11, 163–166.

13. Kharchenko, P.V., Silberstein, L., and Scadden, D.T. (2014). Bayesian approach to single-cell differential expression analysis. Nat Methods 11, 740–742.

14. Kiselev, V.Y., Kirschner, K., Schaub, M.T., Andrews, T., Yiu, A., Chandra, T., Natarajan, K.N., Reik, W., Barahona, M., Green, A.R., et al. (2017). SC3: consensus clustering of single-cell RNA-seq data. Nat Methods 14, 483–486.

15. Law, C.W., Chen, Y., Shi, W., and Smyth, G.K. (2014). voom: Precision weights unlock linear model analysis tools for RNA-seq read counts. Genome Biol 15, R29.

16. Li, Bo, and Colin N. Dewey. 2011. “RSEM: Accurate Transcript Quantification from RNA-Seq Data with or without a Reference Genome.” BMC Bioinformatics 12 (August):

17. Risso, D., Ngai, J., Speed, T.P., and Dudoit, S. (2014). Normalization of RNA-seq data using factor analysis of control genes or samples. Nat Biotechnol 32, 896–902.

18. Ritchie, M.E., Phipson, B., Wu, D., Hu, Y., Law, C.W., Shi, W., and Smyth, G.K. (2015). limma powers differential expression analyses for RNA-sequencing and microarray studies. Nucleic Acids Res 43, e47.

19. Stoeckius, M., Hafemeister, C., Stephenson, W., Houck-Loomis, B., Chattopadhyay, P.K., Swerdlow, H., Satija, R., and Smibert, P. (2017). Simultaneous epitope and transcriptome measurement in single cells. Nat Methods 14, 865–868.

20. Subramanian, A., Tamayo, P., Mootha, V.K., Mukherjee, S., Ebert, B.L., Gillette, M.A., Paulovich, A., Pomeroy, S.L., Golub, T.R., Lander, E.S., et al. (2005). Gene set enrichment analysis: a knowledge-based approach for interpreting genome-wide expression profiles. Proc Natl Acad Sci U S A 102, 15545–15550.

21. Svensson, V., Natarajan, K.N., Ly, L.H., Miragaia, R.J., Labalette, C., Macaulay, I.C., Cvejic, A., and Teichmann, S.A. (2017). Power analysis of single-cell RNA-sequencing experiments. Nat Methods 14, 381–387.

22. Torre, E., Dueck, H., Shaffer, S., Gospocic, J., Gupte, R., Bonasio, R., Kim, J., Murray, J., and Raj, A. (2018). Rare Cell Detection by Single-Cell RNA Sequencing as Guided by Single-Molecule RNA FISH. Cell Syst 6, 171–179 e175.

23. Wang, Y.J., and Kaestner, K.H. (2018). Single-Cell RNA-Seq of the Pancreatic Islets--a Promise Not yet Fulfilled? Cell Metab.

24. Wang, Y.J., Schug, J., Won, K.J., Liu, C., Naji, A., Avrahami, D., Golson, M.L., and Kaestner, K.H. (2016a). Single cell transcriptomics of the human endocrine pancreas. Diabetes.

25. Wang, Y.J., Schug, J., Won, K.J., Liu, C., Naji, A., Avrahami, D., Golson, M.L., and Kaestner, K.H. (2016b). Single-Cell Transcriptomics of the Human Endocrine Pancreas. Diabetes 65, 3028–3038.

26. Wu, A.R., Neff, N.F., Kalisky, T., Dalerba, P., Treutlein, B., Rothenberg, M.E., Mburu, F. M., Mantalas, G.L., Sim, S., Clarke, M.F., et al. (2014). Quantitative assessment of single-cell RNA-sequencing methods. Nat Methods 11, 41–46.

27. Xin, Y., Kim, J., Okamoto, H., Ni, M., Wei, Y., Adler, C., Murphy, A.J., Yancopoulos, G. D., Lin, C., and Gromada, J. (2016). RNA Sequencing of Single Human Islet Cells Reveals Type 2 Diabetes Genes. Cell Metab 24, 608–615.

28. Zhang, X., Li, T., Liu, F., Chen, Y., Yao, J., Li, Z., Huang, Y., and Wang, J. (2019). Comparative Analysis of Droplet-Based Ultra-High-Throughput Single-Cell RNA-Seq Systems. Mol Cell 73, 130–142 e135.

29. Zheng, G.X., Terry, J.M., Belgrader, P., Ryvkin, P., Bent, Z.W., Wilson, R., Ziraldo, S.B., Wheeler, T.D., McDermott, G.P., Zhu, J., et al. (2017). Massively parallel digital transcriptional profiling of single cells. Nature communications 8, 14049.

30. Ziegenhain, C., Vieth, B., Parekh, S., Reinius, B., Guillaumet-Adkins, A., Smets, M., Leonhardt, H., Heyn, H., Hellmann, I., and Enard, W. (2017). Comparative Analysis of Single-Cell RNA Sequencing Methods. Mol Cell 65, 631–643 e634.

